# Analysis-ready VCF at Biobank scale using Zarr

**DOI:** 10.1101/2024.06.11.598241

**Authors:** Eric Czech, Timothy R. Millar, Will Tyler, Tom White, Benjamin Elsworth, Jérémy Guez, Jonny Hancox, Ben Jeffery, Konrad J. Karczewski, Alistair Miles, Sam Tallman, Per Unneberg, Rafal Wojdyla, Shadi Zabad, Jeff Hammerbacher, Jerome Kelleher

## Abstract

**Background:** Variant Call Format (VCF) is the standard file format for interchanging genetic variation data and associated quality control metrics. The usual row-wise encoding of the VCF data model (either as text or packed binary) emphasises efficient retrieval of all data for a given variant, but accessing data on a field or sample basis is inefficient. Biobank scale datasets currently available consist of hundreds of thousands of whole genomes and hundreds of terabytes of compressed VCF. Row-wise data storage is fundamentally unsuitable and a more scalable approach is needed.

**Results:** Zarr is a format for storing multi-dimensional data that is widely used across the sciences, and is ideally suited to massively parallel processing. We present the VCF Zarr specification, an encoding of the VCF data model using Zarr, along with fundamental software infrastructure for efficient and reliable conversion at scale. We show how this format is far more efficient than standard VCF based approaches, and competitive with specialised methods for storing genotype data in terms of compression ratios and single-threaded calculation performance. We present case studies on subsets of three large human datasets (Genomics England: *n*=78,195; Our Future Health: *n*=651,050; All of Us: *n*=245,394) along with whole genome datasets for Norway Spruce (*n*=1,063) and SARS-CoV-2 (*n*=4,484,157). We demonstrate the potential for VCF Zarr to enable a new generation of high-performance and cost-effective applications via illustrative examples using cloud computing and GPUs.

**Conclusions:** Large row-encoded VCF files are a major bottleneck for current research, and storing and processing these files incurs a substantial cost. The VCF Zarr specification, building on widely-used, open-source technologies has the potential to greatly reduce these costs, and may enable a diverse ecosystem of next-generation tools for analysing genetic variation data directly from cloud-based object stores, while maintaining compatibility with existing file-oriented workflows.

**Key Points:**

- VCF is widely supported, and the underlying data model entrenched in bioinformatics pipelines.
- The standard row-wise encoding as text (or binary) is inherently inefficient for large-scale data processing.
- The Zarr format provides an efficient solution, by encoding fields in the VCF separately in chunk-compressed binary format.

## Background

Variant Call Format (VCF) is the standard format for interchanging genetic variation data, encoding information about DNA sequence polymorphisms among a set of samples with associated quality control metrics and metadata [1]. Originally defined specifically as a text file, it has been refined and standardised [2] and the underlying data-model is now deeply embedded in bioinformatics practice. Dataset sizes have grown explosively since the introduction of VCF as part of 1000 Genomes project [3], with Biobank scale initiatives such as Genomics England [4, 5], UK Biobank [6, 7, 8, 9], Our Future Health [10, 11], and the All of Us research program [12] collecting genome sequence data for millions of humans [13]. Large genetic variation datasets are also being generated for other organisms and a variety of purposes including agriculture [14, 15], conservation [16] and infectious disease surveillance [17]. VCF’s simple text-based design and widespread support [18] make it an excellent archival format, but it is an inefficient basis for analysis. Methods that require efficient access to genotype data either require conversion to the PLINK [19, 20] or BGEN [21] formats [e.g. 22, 23, 24] or use bespoke binary formats that support the required access patterns [e.g. 25, 26, 27]. While PLINK and BGEN formats are more efficient to access than VCF, neither can accommodate the full flexibility of the VCF data model and conversion is lossy. PLINK’s approach of storing the genotype matrix in uncompressed packed-binary format provides efficient access to genotype data, but file sizes are substantially larger than the equivalent compressed VCF (see Fig 2). For example, at two bits per diploid genotype, the full genotype matrix for the GraphTyper SNP dataset in the 500K UKB WGS data [9] is 116 TiB.

Processing of Biobank scale datasets can be split into a few broad categories. The most basic analysis is quality control (QC). Variant QC is an involved and multi-faceted task [28, 29, 30, 31], often requiring interactive, exploratory analysis and incurring substantial computation over multiple QC fields. Genotype calls are sometimes refined via statistical methods, for example by phasing [32, 33, 27, 34], and imputation [25, 35, 36, 37] creating additional dataset copies. A common task to perform is a genome wide association study (GWAS) [38]. The majority of tools for performing GWAS and related analyses require data to be in PLINK or BGEN formats [e.g 20, 24, 39, 23], and so data must be “hard-called” according to some QC criteria and exported to additional copies. Finally, variation datasets are often queried in exploratory analyses, to find regions or samples of interest for a particular study [e.g. 40].

VCF cannot support any of these workflows efficiently at the Biobank scale. The most intrinsically limiting aspect of VCF’s design is its row-wise layout of data, which means that (for example) information for a particular sample or field cannot be obtained without retrieving the entire dataset. The file-oriented paradigm is also unsuited to the realities of modern datasets, which are too large to download and often required to stay in-situ by data-access agreements. Large files are currently stored in cloud environments, where the file systems that are required by classical file-oriented tools are expensively emulated on the basic building blocks of object storage. These multiple layers of inefficiencies around processing VCF data at scale in the cloud mean that it is time-consuming and expensive, and these vast datasets are not utilised to their full potential.

To achieve this full potential we need a new generation of tools that operate directly on a primary data representation that supports efficient access across a range of applications, with native support for cloud object storage. Such a representation can be termed “analysis-ready” and “cloud-native” [41]. For the representation to be FAIR [42], it must be *accessible*, using protocols that are “open, free, and universally implementable”. There is currently no efficient, FAIR representation of genetic variation data suitable for cloud deployments. Hail [43, 44] has become the dominant platform for quality control of large-scale variation datasets, and has been instrumental in projects such as gnomadAD [45, 30]. While Hail is built on open components from the Hadoop distributed computing ecosystem [46], the details of its MatrixTable format are not documented or intended for external reuse. Similarly, commercial solutions that have emerged to facilitate the analysis of large-scale genetic variation data are either based on proprietary [47, 48, 49, 50, 51] or single-vendor technologies [e.g. 52, 53]. The next generation of VCF analysis methods requires an open, free and transparent data representation with multiple independent implementations.

In this article, we decouple the VCF data model from its row-oriented file definition, and show how the data can be compactly stored and efficiently analysed in a cloud-native, FAIR manner. We do this by translating VCF data into Zarr format, a method of storing large-scale multidimensional data as a regular grid of compressed chunks. Zarr’s elegant simplicity and first-class support for cloud object stores and parallel computing have led to it gaining substantial traction across the sciences, and it is now used in multiple petabyte-scale datasets in cloud deployments (see Methods for details). We present the VCF Zarr specification that formalises this mapping, along with the vcf2zarr and vcztools utilities to reliably convert VCF to and from Zarr at scale. We show that VCF Zarr is much more compact than VCF and is competitive with state-of-the-art file-based VCF compression tools. Moreover, we show that Zarr’s storage of data in an analysis-ready format greatly facilitates computation, with various benchmarks being substantially faster than bcftools based pipelines, and again competitive with state-of-the-art file-oriented methods. We demonstrate the scalability and flexibility of VCF Zarr via case-studies on five boundary-pushing datasets. We examine conversion and compression performance on three different data modalities for large human datasets: whole genome sequence data from Genomics England (5X smaller than gzipped VCF); genotype data from Our Future Health (2.5X smaller than gzipped VCF); and exome-like data from All of Us (1.1X larger than gzipped VCF). We then examine performance for species with large genomes using Norway Spruce data (3.2X smaller than gzipped VCF), and in a large collection of short whole genome alignments using SARS-CoV-2 data (77X smaller than gzipped FASTA). Interoperability with the vast ecosystem of tools based on VCF [54, 18] is crucial. We first present the vcztools program, which implements a subset of the functionality in bcftools and allows data stored in Zarr at some URL to be treated as if it were a VCF or BCF file, and demonstrate that it can be used as a practical replacement in processing pipelines. We then present an illustrative example of how existing methods can adopt VCF Zarr by modifying the popular SAIGE [55, 56] GWAS software (written in C++), and show significant speedups over the existing VCF (7.9X) and BCF (3.4X) backends in a simple benchmark. Finally, we illustrate the potential for a new generation of Zarr-based applications via some illustative benchmarks. First, we demonstrate the very high levels of throughput performance available on public clouds when data in an object store is accessed in an asynchronous and highly parallel manner. Finally, we demonstrate the ease and efficiency with which data in Zarr format can be processed on a GPU, opening many exciting avenues for future research.

## Results

### Storing genetic variation data

Although VCF is the standard format for exchanging genetic variation data, its limitations both in terms of compression and query/compute performance are well known [e.g. 57, 58, 59], and many methods have been suggested to improve on these properties. Most approaches balance compression with performance on particular types of queries, typically using a command line interface (CLI) and outputting VCF text [58, 59, 60, 61, 62, 63, 64, 65, 66, 67, 68]. Several specialised algorithms for compressing the genotype matrix (i.e., just the genotype calls without additional VCF information) have been proposed [69, 70, 71, 72, 73, 74] most notably the Positional Burrows–Wheeler Transform (PBWT) [75]. (See [76] for a review of the techniques employed in genetic data compression.) The widely-used PLINK binary format stores genotypes in a packed binary representation, supporting only biallelic variants without phase information. The PLINK 2 PGEN format [77] is more general and compact than PLINK, compressing variant data using specialised algorithms [71]. Methods have also been developed which store variation data along with annotations in databases to facilitate efficient queries [e.g. 78, 79] which either limit to certain classes of variant [e.g. 80] or have storage requirements larger than uncompressed VCF [81]. The SeqArray package [82] builds on the Genomic Data Storage container format [83] to store VCF genotype data in a packed and compressed format, and is used in several downstream R packages [e.g. 84, 85].

VCF is a row-wise format in which observations and metadata for a single variant are encoded as a line of text [1]. BCF [86], the standard binary representation of VCF, is similarly row-wise, as are the majority of proposed alternative storage formats. Row-wise storage makes retrieving all information for a given record straight-forward and efficient, and works well when records are either relativelysmall or we typicallywant to analyse each record in its entirety. When we want to analyse only a subset of a record, row-wise storage can be inefficient because we will usually need to retrieve more information than required from storage. In the case of VCF (and BCF) where records are not of a fixed size and are almost always compressed in blocks, accessing any information for a set of rows means retrieving and decompressing *all* information from these rows.

The usual alternative to row-wise storage is *columnar* storage: instead of grouping together all the fields for a record, we group together all the records for a given field. Columnar storage formats such as Parquet [87] make retrieving particular columns much more efficient and can lead to substantially better compression. While columnar techniques have been successfully applied in alignment storage [e.g. 88, 89, 90], columnar technologies for storing and analysing variation data have had limited success [91, 92]. Mapping VCF directly to a columnar layout, in which there is a column for the genotypes (and other per-call QC metrics) for each sample leads to a large number of columns, which can be cumbersome and cause scalability issues. Fundamentally, columnar methods are one-dimensional, storing a vector of values associated with a particular key, whereas genetic variation data is usually modelled as a two-dimensional matrix in which we are interested in accessing both rows *and* columns. Just as row-oriented storage makes accessing data for a given sample inefficient, columnar storage makes accessing all the data for a given variant inefficient.

VCF is at its core an encoding of the genotype matrix, where each entry describes the observed genotypes for a given sample at a given variant site, interleaved with per-variant information and other call-level matrices (e.g., the GQ or AD fields). The data is largely numerical and of fixed dimension, making it a natural fit for array-oriented or “tensor” storage. We propose the VCF Zarr specification which maps the VCF data model into an array-oriented layout using Zarr (Fig 1). In the VCF Zarr specification, each field in a VCF is mapped to a separately-stored array, allowing for efficient retrieval and high levels of compression. See the Methods for more detail on Zarr and the VCF Zarr specification.

**Figure 1.**
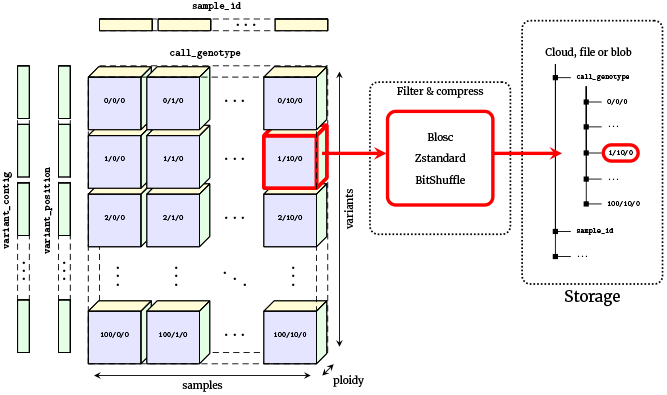
Chunked compressed storage of VCF data using Zarr. The call_genotype array is a three-dimensional (variants, samples, ploidy) array of integers, split into a uniform grid of chunks determined by the variant and sample chunk sizes (10,000 and 1,000 by default in vcf2zarr). Each chunk is associated with a key defining its location in this grid, which can be stored in any key-value store such as a standard file-system or cloud object store. Chunks are compressed independently using standard codecs and pre-compression filters, which can be specified on a per-array basis. Also shown are the one-dimensional variant_contig (CHROM) and variant_position arrays (POS). Other fields are stored in a similar fashion.

One of the key benefits of Zarr is its cloud-native design, but it also works well on standard file systems, where arrays and chunks are stored hierarchically in directories and files (storage as a single Zip archive is also supported, and has certain advantages as discussed later). To enable comparison with the existing file-based ecosystem of tools, we focus on Zarr’s file system chunk storage in a series of illustrative benchmarks in the following sections before presenting cloud-based benchmarks. We compare primarily with VCF/BCF based workflows using bcftools because this is the standard practice, used in the vast majority of cases. We also compare with two representative recent specialised utilities; see [61, 67, 68] for further benchmarks of these and other tools. Genozip [63, 64] is a tool focused on compression performance, which uses a custom file format and a CLI to extract VCF as text with various filtering options. Savvy [65] is an extension of BCF which takes advantage of sparsity in the genotype matrix as well as using PBWT-based approaches for improved compression. Savvy provides a CLI as well as a C++ API. Our benchmarks are based on genotype data from subsets of a large and highly realistic simulation of French-Canadians [93] (see Methods for details on the dataset and bench-marking methodology). Note that while simulations cannot capture all the subtleties of real data, the allele frequency and population structure patterns in this dataset have been shown to closely follow observations [93] and so it provides a reasonable and easily reproducible data point when comparing such methods. The simulations only contain genotypes without any additional high-entropy QC fields, which is unrealistic (see the later case-studies for benchmarks on datasets that contain a wide variety of such fields). Note, however, that such minimal, genotype-only data is something of a best-case scenario for specialised genotype compression methods using row-wise storage.

Fig 2 shows compression performance on up to a million samples for chromosome 21, with the size of the genotype-matrix encoded as 1-bit per haploid call included for reference. Gzip compressed VCF performs remarkably well, compressing the data to around 5% of the minimal binary encoding of a biallelic genotype matrix for 1 million samples. BCF provides a significant improvement in compression performance over VCF (note the log-log scale). Genozip has superb compression, having far smaller file sizes that the other methods (although somewhat losing its advantage at larger sample sizes). Zarr and Savvy have very similar compression performance in this example (23.75 GiB and 21.25 GiB for 10^6^ samples, respectively). When using a chunk size of 10^4^ variants × 10^3^ samples, Zarr storage is reduced to 22.07 GiB for 10^6^ samples (data not shown; see Figs S9 and S10 for more analysis on the effect of chunk size on compression). It is remarkable that the simple approach of compressing two dimensional chunks of the genotype matrix using the Zstandard compressor [94] and the bit-shuffle filter from Blosc [95] (see Methods for details) produces compression levels competitive with the highly specialised methods used by Savvy.

**Figure 2.**
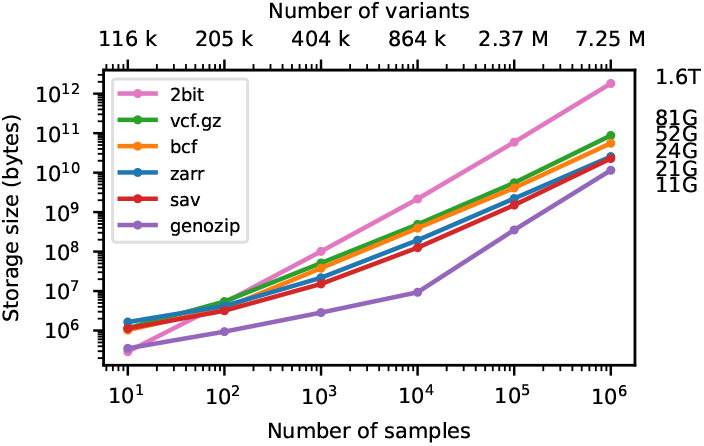
Compression performance on simulated genotypes. Comparison of total stored bytes for VCF data produced by subsets of a large simulation of French-Canadians. Sizes for 10^6^ samples are shown on the right. Also shown for reference is the size of genotype matrix when encoded as two bits per diploid genotype (2bit), as used in the PLINK binary format. Default compression settings were used in all cases.

### Calculating with the genotype matrix

Storing genetic variation data compactly is important, but it is also important that we can analyse the data efficiently. Bioinformatics workflows tend to emphasise text files and command line utilities that consume and produce text [e.g. 96]. Thus, many tools that compress VCF data provide a command line utility with a query language to restrict the records examined, perform some pre-specified calculations and finally output some text, typically VCF or tab/comma separated values [58, 59, 61, 62, 63, 64, 67, 68]. These pre-defined calculations are by necessity limited in scope, however, and the volumes of text involved in Biobank scale datasets make the classical approach of custom analyses via Unix utilities in pipelines prohibitively slow. Thus, methods have begun to provide Application Programming Interfaces (APIs), providing efficient access to genotype and other VCF data [e.g. 57, 65, 66]. By providing programmatic access, the data can be retrieved from storage, decoded and then analysed in the same memory space without additional copies and inter-process communication through pipes.

To demonstrate the accessibility of genotype data and efficiency with which calculations can be performed under the different formats, we use the bcftools +af-dist plugin (which computes a table of deviations from Hardy-Weinberg expectations in allele frequency bins) as an example. We chose this particular operation for several reasons. First, it is a straightforward calculation that requires examining every element in the genotype matrix, and can be reproduced in different programming languages without too much effort. Secondly, it produces a small volume of output and therefore the time spent outputting results is negligible. Finally, it has an efficient implementation written using the htslib C API [97], and therefore running this command on a VCF or BCF file provides a reasonable approximation of the limit of what can be achieved in terms of whole-matrix computation on these formats.

Fig 3 shows timing results for running bcftools +af-dist and equivalent operations on the data of Fig 2. There is a large difference in the time required (note the log-log scale). The slowest approach uses Genozip. Because Genozip does not provide an API and only outputs VCF text, the best approach available is to pipe its output into bcftools +af-dist. This involves first decoding the data from Genozip format, then generating large volumes of VCF text (terabytes, in the largest examples here), which we must subsequently parse before finally doing the actual calculation. Running bcftools +af-dist directly on the gzipped VCF is substantially faster, indicating that Genozip’s excellent compression performance comes at a substantial decompression cost. Using a BCF file is again significantly faster, because the packed binary format avoids the overhead of parsing VCF text into htslib’s internal data structures. We only use BCF for subsequent bcftools benchmarks.

**Figure 3.**
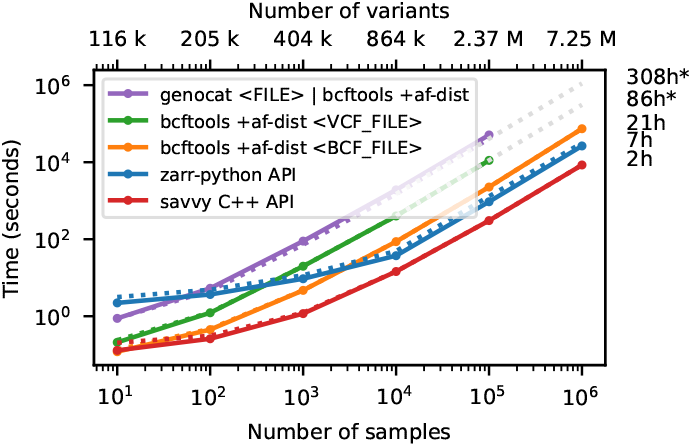
Whole-matrix compute performance with increasing sample size. Total CPU time required to run bcftools +af-dist and equivalent operations in a single thread for various tools. Elapsed time is also reported (dotted line). Run-time for genozip and bcftools on VCF at 10^6^ samples were extrapolated by fitting an exponential. See Methods for full details.

The data shown in Fig 3 for Zarr and Savvy is based on custom programs written using their respective APIs to implement the af-dist operation. The Zarr program uses the Zarr-Python package to iterate over the decoded chunks of the genotype matrix and classifies genotypes within a chunk using a 14 line Python function, accelerated using the Numba JIT compiler [98]. The allele frequencies and genotype counts are then analysed to produce the final counts within the allele frequency bins with 9 lines of Python using NumPy [99] functions. Remarkably, this short and simple Python program is substantially faster than the equivalent compiled C using htslib APIs on BCF (7.3 hours vs 20.6 hours for 1 million samples). The fastest method is the C++ program written using the Savvy API. This would largely seem to be due to Savvy’s excellent genotype decoding performance (up to 6.6 GiB/s vs 1.2 GiB/s for Zarr on this dataset; Fig S1). Turning off the BitShuffle filter for the Zarr dataset, however, leads to a substantial increase in decoding speed (2.7 GiB/s) at the cost of a roughly 25% increase in storage space (31.2 GiB up from 23.75 GiB for 1 million samples; data not shown). The LZ4 codec (again without BitShuffle) leads to slightly higher decoding speed (2.9 GiB/s), but at the cost of much larger storage requirements (214 GiB; data not shown). Given the relatively small contribution of genotypes to the overall storage of real datasets (see the Genomics England example) and the frequency that they are likely to be accessed, using Zstandard without BitShuffle would seem like a good tradeoff in many cases. This ability to easily tune compression performance and decoding speed on a field-by-field basis is a major strong point of Zarr. The vcf2zarr utility also provides functionality to aid with such storage schema tuning.

The discrepancy between elapsed time and total CPU time for Zarr benchmarks in Figs 3– 5 and S1 is mostly due to the overhead of accessing a large number of small chunk files and consequent reduced effectiveness of operating system read-ahead caching. While having many independent chunks maps naturally to high-performance parallel access patterns in the cloud store setting [100] (see also the section on accessing Genomics England data stored on S3), such file access patterns can be problematic in the classical file-system setting. The Zarr Zip store (which is simply a Zip archive of the standard directory hierarchy) provides a useful alternative here and can significantly reduce overall processing time. (The nascent Zarr v3 specification provides another option via “sharding” which essentially allows multiple chunks to be stored in a single file.) To illustrate this we repeated the benchmarks in Fig S1 in which we measure the time required to decode the entire genotype matrix (thus serving as a lower-bound for any subsequent computations). In the case of 10^6^ samples, the Zip file requires slightly more storage space than the directory hierarchy (23.75 GiB and 24.24 GiB for file system and Zip file, respectively) and while total CPU time is similar (193m vs 187m), the elapsed wall clock time for the Zip file version is significantly less (258m vs 188m).

Zarr is a standard with multiple independent implementations across different programming languages (see Methods). Fig S2 compares the CPU time required to perform the computations of Fig 3 for af-dist implementations using the Zarr-Python [101], JZarr [102], and TensorStore [103] (C++ and Python) libraries. The program using the TensorStore C++ API is fastest, requiring about 20% less CPU time than the version using Zarr-Python and Numba used in our other benchmarks.

### Subsetting the genotype matrix

As datasets grow ever larger, the ability to efficiently access subsets of the data becomes increasingly important. VCF/BCF achieve efficient access to the data for genomic ranges by compressing blocks of adjacent records using bgzip, and storing secondary indexes alongside the original files with a conventional suffix [104]. Thus, for a given range query we decompress only the necessary blocks and can quickly access the required records. The row-wise nature of VCF (and most proposed alternatives), however, means that we cannot efficiently subset *by sample* (e.g., to calculate statistics within a particular cohort). In the extreme case, if we want to access only the genotypes for a single sample we must still retrieve and decompress the entire dataset.

We illustrate this cost of row-wise encoding in Fig 4, where we run the af-dist calculation on a small fixed-size subset of the genotype matrices of Fig 2. The two-dimensional chunking of Zarr means that this sub-matrix can be efficiently extracted, and therefore the execution time depends very weakly on the overall dataset size, with the computation requiring around 2 seconds for 1 million samples. Because of their row-wise encoding, CPU time scales with the number of samples for all the other methods. Fig S3 shows performance for the same operation when selecting half of the samples in the dataset.

**Figure 4.**
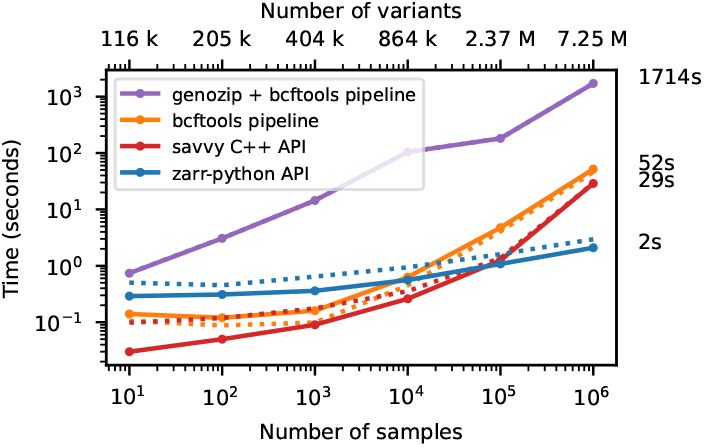
Compute performance on subsets of the matrix. Total CPU time required to run the af-dist calculation for a contiguous subset of 10,000 variants × 10 samples from the middle of the matrix for the data in Fig 2. Elapsed time is also reported (dotted line). The genozip and bcftools pipelines involve multiple commands required to correctly calculate the AF INFO field required by bcftools +af-dist. See the Methods for full details on the steps performed.

### Extracting, inserting and updating fields

We have focused on the genotype matrix up to this point, contrasting Zarr with existing row-wise methods. Real-world VCFs encapsulate much more than just the genotype matrix, and can contain large numbers of additional fields. Fig 5 shows the time required to extract the genomic position of each variant in the simulated benchmark dataset, which we can use as an indicative example of a per-variant query. Although Savvy is many times faster than bcftools query here, the row-wise storage strategy that they share means that the entire dataset must be read into memory and decompressed to extract just one field from each record. Zarr excels at these tasks: we only read and decompress the information required.

**Figure 5.**
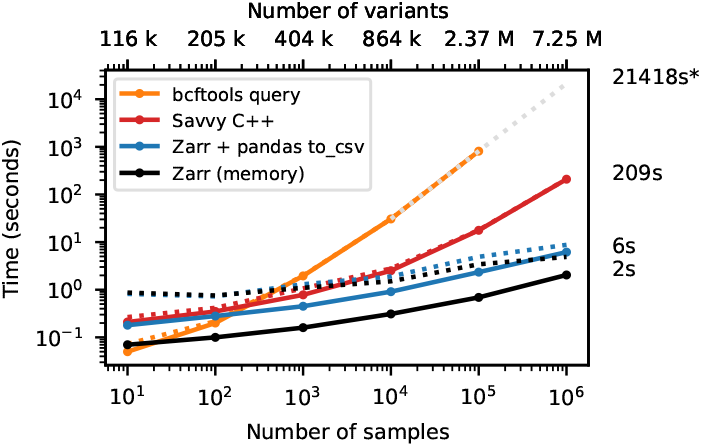
Time to extract the genome position and write to a text file. Total CPU time required to extract the POS field for BCF, sav and Zarr formats for the data in Figure 2. For the BCF file we used bcftools query -f”%POS\n”. For sav, we used the Savvy C++ API to extract position for each variant and output text using the std::cout stream. For Zarr, we read the variant_position array into a NumPy array, and then wrote to a text file using the Pandas write_csv method. Zarr CPU time is dominated by writing the text output; we also show the time required to populate a NumPy array with the data in Zarr, which is 2 seconds. Wall-clock time (dotted line) is dominated in this case by file I/O. Note that our benchmarking methodology of dropping the file system caches before each run (see Methods) is likely not reflective of real world usage where a small and frequently used field such as POS would remain in cache. Time to output text for Savvy is not significant for > 1000 samples (not shown).

Many of the additional fields that we find in real-world VCFs are variant-level annotations, extensively used in downstream applications. For example, a common workflow is to add or update variant IDs in a VCF using a reference database such as dbSNP [105]. The standard approach to this (using e.g. bcftools annotate) is to create a *copy* of the VCF which includes these new annotations. Thus, even though we may only be altering a single field comprising a tiny fraction of the data, we still read, decompress, update, compress and write the entire dataset to a new file. With Zarr, we can update an existing field or add arbitrary additional fields without touching the rest of the data or creating redundant copies.

### Case study: Genomics England WGS data

In this section we demonstrate the utility of VCF Zarr on a large human dataset and the scalability of the vcf2zarr conversion utility. Genomics England’s multi-sample VCF dataset (aggV2) is an aggregate of 78,195 gVCFs from rare disease and cancer participants recruited as part of the 100,000 Genomes Project [4, 5]. The dataset comprises approximately 722 million annotated single-nucleotide variants and small indels split into 1,371 roughly equal chunks and totalling 165.3 TiB of VCF data after bgzip compression. The dataset is used for a variety of research purposes, ranging from GWAS [106] and imputation [107] to simple queries involving single gene regions [108, 109].

As described in the Methods, conversion to Zarr using vcf2zarr is a two-step process. We first converted the 106 VCF files (12.81 TiB) for chromosome 2 into the intermediate columnar format (ICF). This task was split into 14,605 partitions, and distributed using the Genomics England HPC cluster. The average run-time per partition was 20.7 min. The ICF representation used a total of 9.94 TiB over 3,960,177 data storage files. We then converted the ICF to Zarr, partitioned into 5989 independent jobs, with an 18.6 min average run time. This produced a dataset with 44 arrays, consuming a total of 2.54 TiB of storage over 6,312,488 chunk files (using a chunk size of 10^4^ variants × 10^3^ samples). This is a roughly 5X reduction in total storage space over the original VCF. The top fields in terms of storage are detailed in Table 1. We do not compare with other tools such as Genozip and Savvy here because they have fundamental limitations (as shown in earlier simulation-based benchmarks), and conversion of these large VCFs is a major undertaking.

**Table 1.** Summary for VCF Zarr conversion of Genomics England WGS data for chromosome 2 (78,195 samples, 59,880,903 variants), consisting of 44 fields and 2.54 TiB of storage (∼5X compression over source gzipped VCF). Each field is stored independently as a Zarr array with the given type (sufficient to represent all values in the data). We show the total storage consumed (reported via du) in power-of-two units, and the compression ratio achieved on that array. We also show the percentage of the overall storage that each array consumes. Shown are the top 11 fields consuming at least 0.01% of the overall storage.

| Field | Type | Storage | Compress | %Total |
| --- | --- | --- | --- | --- |
| `call_AD` | int16 | 658.44GiB | 26.0 | 25.35 % |
| `call_GQ` | int16 | 654.45GiB | 13.0 | 25.20 % |
| `call_DP` | int16 | 570.03GiB | 15.0 | 21.95 % |
| `call_DPF` | int16 | 447.09GiB | 20.0 | 17.22 % |
| `call_PL` | int16 | 162.56GiB | 160.0 | 6.26 % |
| `call_GQX` | int16 | 40.99GiB | 210.0 | 1.58 % |
| `call_FT` | str | 25.04GiB | 1400.0 | 0.96 % |
| `call_genotype` | int8 | 21.46GiB | 410.0 | 0.83 % |
| `call_genotype_mask` | bool | 12.78 GiB | 680.0 | 0.49 % |
| `call_genotype_phased` | bool | 2.35 GiB | 1900.0 | 0.09 % |
| `call_PS` | int8 | 383.38 MiB | 12 000.0 | 0.01 % |
| `call_ADF` | int8 | 383.38 MiB | 12 000.0 | 0.01 % |
| `call_ADR` | int8 | 383.38 MiB | 12 000.0 | 0.01 % |

Table 1 shows that the dataset storage size is dominated by a few columns with the top four (call_AD, call_GQ, call_DP and call_DPF) accounting for 90% of the total. These fields are much less compressible than genotype data (which uses < 1% of the total space here) because of their inherent noisiness [62]. Note that these top four fields are stored as 16 bit integers because they contain rare outliers that cannot be stored as 8 bits. While the fields could likely be truncated to have a maximum of 127 with minimal loss of information, the compression gains from doing so are relatively minor, and we therefore opt for fully lossless compression here for simplicity. The call_PS field here has an extremely high compression ratio because it consists entirely of missing data (i.e., it was listed in the header but never used in the VCF).

To demonstrate the computational accessibility of Zarr on this large human dataset, we performed some illustrative benchmarks. As these benchmarks take some time to run, we focus on a single 132 GiB compressed VCF file covering positions 58,219,159–60,650,943 (562,640 variants) from the middle of the list of 106 files for chromosome 2. We report both the total CPU time and elapsed wall-clock time here as both are relevant. First, we extracted the genome position for each variant in this single VCF chunk using bcftools query and Python Zarr code as described in Fig 5. The bcftools command required 55.42 min CPU and 85.85 min elapsed. The Zarr code required 2.78 sec CPU and 1.73 min elapsed. This is a 1196X smaller CPU burden and a 50X speed-up in elapsed time. The major difference between CPU time and wall-time is noteworthy here, and highlights the importance of using an index to find the location of variant chunks. This benchmark predates the region_index array computed by vcf2zarr (see Methods) which stores sufficient information to identify which variant chunks intersect with a given range query. Here, to find the relevant slice of records corresponding to the range of positions in the target VCF file, we simply load the entire variant_position array and binary search. This entails reading 5,989 chunk files (the chunk size is 100,000 variants) and incurred a substantial latency penalty on the Genomics England HPC shared filesystem.

We then ran the af-dist calculation (Figs 3 and 4) on the VCF file using bcftools +af-dist as before. The elapsed time for this operation was 716.28 min CPU, 716.3 min elapsed. Repeating this operation for the same coordinates in Zarr (using Python code described in previous sections) gave a total CPU time of 2.32 min and elapsed time of 4.25 min. This is a 309X reduction in CPU burden and a 169X speed-up in elapsed time. It is worth noting here that bcftools +af-dist cannot be performed in parallel across multiple slices of a chromosome, and if we did want to run it on all of chromosome 2 we would need to concatenate the 106 VCF files. While af-dist itself is not a common operation, many tasks share this property of not being straightforwardly decomposable across multiple VCF files.

Additionally, to illustrate performance on a common filtering task, we created a copy of the VCF chunk which contains only variants that pass some common filtering criteria using bcftools view -I –include “FORMAT/DP>10 & FORMAT/GQ>20”, following standard practices [e.g. 110, 106, 30]. This used 689.46 min CPU time, with an elapsed time of 689.48 min. In comparison, computing and storing a variant mask (i.e., a boolean value for each variant denoting whether it should be considered or not for analysis) based on the same criteria using Zarr consumed 1.96 min CPU time with an elapsed time of 11 min. This is a 358X reduction in CPU usage, and 63X reduction in elapsed time. There is an important distinction here between creating a copy of the data (an implicit part of VCF based workflows) and creating an additional *mask*. As Table 1 illustrates, call-level masks are cheap (the standard genotype missingness mask, call_genotype_mask, uses 0.49% of the overall storage) and variant or sample level masks require negligible storage. If downstream software can use configurable masks (at variant, sample and call level) rather than expecting full copies of the data, major storage savings and improvements in processing efficiency can be made. The transition from the manifold inefficiencies of present-day “copy-oriented” computing, to the “mask-oriented” analysis of large immutable, single-source datasets is a potentially transformational change enabled by Zarr.

### Case study: Our Future Health genotype data

Our Future Health [10, 11] is the UK’s largest ever health research programme. It is designed to help people live healthier lives for longer through the discovery and testing of more effective approaches to prevention, earlier detection and treatment of diseases. Data Release 9 [111] was made available in December 2024, including genotype array data for from 651,050 individuals and 707,522 variants. For this study, we converted the genotype data for chromosome 22 consisting of 10,221 variants over three gzipped VCF files and a total of 42.3 GiB of storage.

Conversion was performed on a Standard_D32s_v5 Azure VM instance, with 32 CPUs and 128 GB RAM. We first converted the VCF data to ICF, which took 8m 55s elapsed (138m CPU) using 20 processes. The ICF representation required a total of 41 GiB of storage. We then converted to VCF Zarr format using the default schema, which resulted in a 41 GiB VCZ over 4514 chunk files. On inspection, this was dominated by three floating point fields LRR (Log R Ratio), BAF (B allele frequency) and GS (GenCall Score) stored to a high degree of precision (7 decimal places for LRR and BAF and 4 decimal places for GS). On inspecting the histograms for these fields (Fig S6) it seemed that such high precision was likely unnecessary, and we investigated applying the Quantize and BitRound filters from numcodecs [112] to reduce the precision in stored floating point data, and therefore improve compression. Using the Quantize filter with 5 digits of precision on these three fields reduced the overall storage space to 31 GiB with a mean absolute error (MAE, i.e., the average difference between the truncated and original values) of 1.91 × 10^−6^ (LRR), 1.18 × 10^−6^ (BAF) and 1.88 × 10^−6^ (GS). Using the BitRound filter with a parameter of 5 bits reduced the overall storage for the Zarr to 17 GiB with an MAE of 6.93 × 10^−4^ (LRR), 7.86 × 10^−4^ (BAF) and 3.38 × 10^−3^ (GS). The histograms of the truncated fields in all cases were very similar to the to the original (Fig S6). Table 2 shows a summary of the Zarr dataset using the BitRound filter.

**Table 2.** Summary for VCF Zarr conversion of Our Future Health genotype data for chromosome 22 (651,050 samples, 10,221 variants) consisting of 19 fields and 17 GiB of storage (∼ 2.5X compression over source gzipped VCF). Here we use a chunk size of 1,000 variants × 10,000 samples and apply the BitRound(5) filter to the large floating point fields (see text for details). Shown are the top 6 fields consuming at least 0.01% of the overall storage (see Table 1 for column details).

| Field | Type | Storage | Compress | % Total |
| --- | --- | --- | --- | --- |
| call_LRR | float32 | 8.35 GiB | 3.0 | 52.08% |
| call_BAF | float32 | 6.02 GiB | 4.1 | 37.55% |
| call_GS | float32 | 961.37 MiB | 26.0 | 5.86% |
| call_genotype | int8 | 657.21 MiB | 19.0 | 4.00% |
| call_genotype_mask | bool | 79.48 MiB | 160.0 | 0.48% |
| sample_id | str | 2.93 MiB | 1.7 | 0.02% |

### Case study: All of Us exome-like data

The All of Us dataset comprises 245K clinical-grade whole genome sequences, identifying over 1 billion variants, including 3.9 million novel variants with coding consequences [12]. Participants are from the United States, with an emphasis on historically underrepresented genetic ancestry groups. The genomic data, generated through short-read sequencing on the Illumina NovaSeq 6000, is of high quality, with an average coverage depth of 30x, and is linked to extensive health-related information. For this study, we converted the chromosome 20 exome-like data from the All of Us dataset to VCF Zarr. The exome-like data is constructed by extracting from all WGS samples the SNP and indel variants that lie within the chromosome 20 exon regions of the Gencode v42 basic transcripts. This dataset, stored as a single compressed VCF file of 7.44 GiB, includes 715,256 variants, of which more than 700 are classified as pathogenic or likely pathogenic.

Conversion was performed on the All of Us Researcher Workbench platform (based on Google Cloud), using a single compute node with 64 CPUs and 416 GiB of RAM, which cost $3.80/hour. We first converted the VCF to ICF, which took 6 hours and 17 minutes of effective runtime and used 343 hours of CPU time, achieving a parallel efficiency of 86%. This step incurred a cost of $23.88. The resulting ICF representation required 7.5 GiB of storage distributed across 91,232 files. Next, the ICF was converted to Zarr, which took 1 hour and 8 minutes, with a CPU time of 68 hours and 8 minutes and a parallel efficiency of 94%, at a cost of $4.31. This second Zarr representation occupied 8.5 GiB of storage across 232,809 chunk files.

Local alleles (see Methods) is essential in this example as the maximum number of alleles observed at a site is 95. Thus, as defined, the AD field field has an inner dimension of 95 each 1000 × 10000 chunk has an uncompressed size of 1.77 GiB (in this 16 bit integers are required to store the values). Given that the vast majority of this encoding consists of the dimension padding sentinel (−2, see Methods), it is an inefficient means of representing and processing allele depth data. Translating to the local alleles representation here reduces the inner dimension to 2, and the uncompressed size of each chunk to 38.15 MiB. Note that there is also no loss of information in this case by encoding as local alleles, as the per-sample gVCFs used in earlier stages in the All of Us pipeline were truncated to a maximum of two ALT alleles per site.

At 1.1X larger than the source gzipped VCF, the relatively poor compression performance here is noteworthy and interesting. At only 7.44 GiB for for 245K samples and 715K variants, VCF compression performance is exceptional and a result of several data reduction techniques. Firstly, data for the AD, RGQ and FT fields is not stored for homozygous reference calls and the fields are consequently extremely sparse. This is reflected in the Zarr, where compression performance for e.g. LAD is more than 10X higher than the AD field in the Genomics England data (Table 1). Secondly, the GQ field has a very limited range of values as a result of gVCF reblocking. Together, these techniques mean that the VCF text stream is highly compressible. While the Zarr does not compress as well as VCF text in this case, the sparsity of the QC columns is reflected in the very high compression rates. It is interesting to note that the BCF representation of this file is also larger than the VCF (and Zarr) at 8.8GiB.

### Case study: 1,063 spruce whole-genome samples

To demonstrate the versatility of vcf2zarr, in this section we include a case study of a dataset originating from a species whose genome properties differ substantially from human. The conifer Norway spruce, *Picea abies*, is one of the largest and most ecologically important species on Earth and is distributed over large parts of the Northern hemisphere. One of its features is its large genome, consisting of 12 autosomes with a genome size of 19.6 Gb [113]. Here, we assess the performance of vcf2zarr as applied to 1,063 whole-genome resequenced spruce individuals from a recent study (in preparation), where resequencing data has been mapped to a chromosome-scale reference genome. The dataset consists of approximately 3.75 billion single-nucleotide variants and small indels from the 12 autosomes totalling 7.4TiB of VCF data after bgzip compression, distributed over 165 VCF files.

We first converted the 165 VCF files (7.33 TiB) to ICF, followed by conversion to Zarr. The tasks were run as single jobs on a compute node with 512 GB RAM and 128 cores using 120 cores. Conversion to ICF required 13h45min runtime. The ICF representation used a total of 6.77 TiB over 1,340,804 data storage files. Conversion to Zarr using the default schema and a chunk size of 10,000 variants × 1,063 samples required 18h36min runtime, generating a dataset with 34 arrays, and required a total of 6.6 TiB storage over 10,861,038 chunk files. Over 90% of this was used by the call_PL field, mostly due to the maximum number of alleles at a site being 4, and consequently the inner dimension of call_PL 10 because of its quadratic dependency on the number of alleles (see Methods).

We therefore generated a vcf2zarr schema using local allele fields with vcf2zarr mkschema –local-alleles, and used the generated schema as input to vcf2zarr encode. Zarr conversion required 14h09min runtime, generating a dataset with 35 arrays, consuming a total of 2.3 TiB storage over 11,235,556 chunk files. Using local allele fields in this example we therefore achieve approximately 3X improvement of compression. The top fields in terms of storage are detailed in Table 4.

**Table 3.** Summary for VCF Zarr conversion of All of Us exome-like data genotype data for chromosome 20 (245,394 samples, 715,256 variants) consisting of 25 fields and 8.5 GiB of storage (∼ 1.1X larger than source gzipped VCF; see text for discussion). Here we use the local alleles fields call_LA and call_LAD; see text for details. Shown are the top 14 fields consuming at least 0.01% of the overall storage (see Table 1 for column details). The variant_homozygote_count field has been renamed variant_hom_count for display purposes.

| Field | Type | Storage | Compress | % Total |
| --- | --- | --- | --- | --- |
| call_LAD | int16 | 2.26 GiB | 290.0 | 27.85% |
| call_GQ | int8 | 1.89 GiB | 87.0 | 23.29% |
| call_LA | int8 | 1.24 GiB | 260.0 | 15.28% |
| call_genotype | int8 | 914.22 MiB | 370.0 | 11.00% |
| call_RGQ | int16 | 729.57 MiB | 460.0 | 8.78% |
| call_genotype_mask | bool | 606.02 MiB | 550.0 | 7.29% |
| call_FT | str | 498.61 MiB | 2700.0 | 6.00% |
| call_genotype_phased | bool | 17.17 MiB | 9700.0 | 0.21% |
| variant_AF | float32 | 5.95 MiB | 43.0 | 0.07% |
| variant_hom_count | int32 | 4.78 MiB | 54.0 | 0.06% |
| variant_allele | str | 4.74 MiB | 110.0 | 0.06% |
| variant_AC | int32 | 4.24 MiB | 61.0 | 0.05% |
| variant_filter | bool | 2.87 MiB | 0.9 | 0.03% |
| variant_AN | int32 | 904.75 KiB | 3.1 | 0.01% |

**Table 4.** Summary for VCF Zarr conversion of the Norway Spruce WGS data on all 12 autosomes (1,063 samples, 3,745,170,452 variants) consisting of 34 fields and 2.3 TiB of storage (∼ 3.2X smaller than source gzipped VCFs). The chunk size is 10,000 variants × 1,063 samples (i.e., a single sample chunk). This uses local alleles to reduce the space required by the PL field; see the text for discussion. Shown are the top 21 fields consuming at least 0.01% of the overall storage (see Table 1 for column details).

| Field | Type | Storage | Compress | % Total |
| --- | --- | --- | --- | --- |
| call_LPL | int16 | 1.26TiB | 17.0 | 54.76 % |
| call_LA | int8 | 415.39 GiB | 18.0 | 17.63 % |
| call_genotype | int8 | 381.28 GiB | 19.0 | 16.18 % |
| call_genotype_mask | bool | 103.84 GiB | 71.0 | 4.41 % |
| variant_DP4 | int32 | 26.24 GiB | 2.1 | 1.11 % |
| variant_MQSBZ | float32 | 13.08 GiB | 1.1 | 0.56 % |
| variant_RPBZ | float32 | 13.04 GiB | 1.1 | 0.55 % |
| variant_MQBZ | float32 | 12.97 GiB | 1.1 | 0.55 % |
| variant_SCBZ | float32 | 12.94 GiB | 1.1 | 0.55 % |
| variant_quality | float32 | 12.84 GiB | 1.1 | 0.55 % |
| variant_VDB | float32 | 12.84 GiB | 1.1 | 0.55 % |
| variant_BQBZ | float32 | 12.64 GiB | 1.1 | 0.54 % |
| variant_SGB | float32 | 12.12 GiB | 1.2 | 0.51 % |
| variant_position | int32 | 9.65 GiB | 1.4 | 0.41 % |
| variant_AC | int16 | 7.42 GiB | 2.8 | 0.31 % |
| variant_DP | int32 | 6.71 GiB | 2.1 | 0.28 % |
| variant_allele | str | 5.37 GiB | 21.0 | 0.23 % |
| variant_AN | int16 | 3.05 GiB | 2.3 | 0.13 % |
| call_genotype_phased | bool | 1.75 GiB | 2100.0 | 0.07 % |
| variant_filter | bool | 1.46 GiB | 2.4 | 0.06 % |
| variant_MQ | int8 | 489.41 MiB | 7.3 | 0.02 % |

Large genomes from non-model organisms pose particular problems for analysis workflows that often are tailored for human conditions. For instance, the binary Sequence Alignment/Map index format BAI [114] has a 512 Mbp limit, but spruce chromosomes range in size from 1.0 to 1.7 Gbp. The spruce BAM files used for variant calling were BAI indexed, which motivated the partitioning into multiple VCF files in the first place, adding complexity to downstream processing. A similar limitation is imposed by the VCF POS field which is encoded in 32-bit format and can hold chromomosomes up to 2.14 Gbp. Although spruce does not have chromosomes that exceed the limit, other species do, such as the 87.2 Gbp South American lungfish, whose 19 chromosomes are all but one larger than the 3.1 Gbp human genome [115]. Zarr’s flexible type system means that coordinates can be stored as 64 or 32 bit integers as needed, and there is therefore no particular limit on genome size.

### Case study: SARS-CoV-2 alignment data

The SARS-CoV-2 pandemic resulted in viral sequence data collection at an unprecedented scale, with over 15 million whole genomes incorporated into one resource as of 2024 [116]. Data at such scale overwhelmed classical phylogenetic methods, leading to the development of a new class of tools [117, 118, 119] capable of handling millions of samples. The Viridian dataset [120] is a consistently assembled set of consensus sequences for 4,484,157 samples, addressing a range of systematic errors. The sc2ts inference tool uses VCF-Zarr as its input format, and Zhan et al [119] have made the Viridian dataset, aligned to the reference using MAFFT [121], available for download [122]. This dataset has quite different properties to those examined in previous case studies, and illustrates the flexibility of Zarr across a range of biological applications.

One of the unusual properties of this dataset is that whole genome alignments are stored, such that information on all 29,903 bases is stored for every sample. Given the number of samples and that nearly all bases have variation (all sites have at least one sample with missing data, and 99.4% have more than one allele) it is simply more convenient to store the entire alignment for each sample than to manage the complexity of subsetting the sites. Similarly, the usual convention in VCF Zarr is that the variant_alleles field contains the REF and ALT alleles as observed (such that the zero’th allele is always the reference). In this case it is more convenient to encode the alleles as the original bases and IUPAC uncertainty codes [123], such that allele 0 is always A, 1 is C etc. Other than these minor deviations in interpretation, the Zarr follows the VCF Zarr specification and can be processed using the same tools.

Decompressed, the Viridian dataset consists of 48 FASTA files totalling 125 GiB of storage and a 1.4 GiB TSV file for sample metadata. When aligned to the reference and converted to Zarr, the entire dataset is stored in a single 401 MiB Zip file. When compressed with bgzip (so they can be indexed; see next paragraph) the alignments require 22 GiB of storage. Table 5 summarises the storage used by individual arrays, and shows that the alignment data requires 279 MiB, around 77X smaller than gzipped FASTA. The original dataset is distributed as xz compressed FASTA files, which achieves even higher compression levels at a total of 218 MiB. However, these files must be decompressed in order to access the data using standard FASTA processing techniques.

**Table 5.** Summary for VCF Zarr conversion of Viridian SARS-CoV-2 whole genome alignments and metadata (4,484,157 samples, 29,903 variants) consisting of 37 fields and 401 MiB of storage, of which 279 is used by alignments (∼ 77X smaller than gzipped FASTA). The chunk size is 100 variants × 10,000 samples. Shown are the top 9 fields consuming at least 1% of the overall storage (see Table 1 for column details). Field names (derived from source TSV file) are shortened for display purposes by replacing “Genbank” with “G” and “Viridian” with “V”.

| Field | Type | Storage | Compress | % Total |
| --- | --- | --- | --- | --- |
| call_genotype | int8 | 279.38 MiB | 447.5 | 80.04 % |
| sample_G_tree_name | str | 13.18 MiB | 2.5 | 3.78 % |
| sample_G_accession | str | 7.31 MiB | 4.6 | 2.09 % |
| sample_Sample | str | 7.26 MiB | 4.6 | 2.08 % |
| sample_Experiment | str | 6.9 MiB | 4.8 | 1.98 % |
| sample_id | str | 6.83 MiB | 4.9 | 1.96 % |
| sample_V_N | int16 | 4.32 MiB | 1.9 | 1.24 % |
| sample_V_pangolin_1.29 | str | 3.98 MiB | 8.4 | 1.14 % |
| sample_V_pangolin | str | 3.89 MiB | 8.6 | 1.11 % |

Computational accessibility for the Zarr is excellent. For example, computing the total number of missing or ambiguous bases for all 4.4 million samples required 57.4 seconds elapsed (7min 08s CPU) using 8 threads. This required 11 lines of code, using only the Python standard library and NumPy. Similarly, computing allele counts at each site required 4min 19s (20min 50s CPU) using a slightly more complicated 16 line Python function. Existing methods for working with FASTA data provided efficient access to individual alignments using the fai index format, which works with uncompressed or bgzip compressed files. For example, retrieving the alignment for the 1 millionth sample from the corresponding chunk file requires only a few milliseconds using pyfaidx [124] or pysam [125]. Retrieving the same alignment using Zarr required 327ms. Accessing calls for all samples at at a given site requires a similar time (here, accessing site 14,951 required 482ms). FASTA cannot support efficient retrieval of data for a particular site in a set of alignments. Storing data chunked in two dimensions means that we can avoid the classic choice between sample-major or variant-major storage, and access it efficiently in both dimensions.

### Interoperability: vcztools

To provide compatibility with existing workflows we have developed the vcztools command line utility which implements a subset of bcftools functionality. Vcztools is intended to be a drop-in replacement for bcftools, providing an evolutionary path for VCF Zarr adoption, with compatibility being a core objective. Vcztools currently supports Zarr stored in the local file system, Zip archives, and major cloud stores. Thus, users can convert their VCF data to Zarr using bio2zarr, store it in a local file system, Zip file or cloud store as desired and then access this data source within existing processing pipelines, essentially replacing bcftools view <vcf file> with vcztools view <Zarr URL>. This allows established analysis workflows to coexist with new Zarr-native approaches, working from the same primary data.

The initial version of vcztools implements a subset of the query and view commands. We provide an example based on the simulated dataset of 10^5^ samples of Fig 2 to illustrate the functionality and performance of vcztools. First, we extract the position column using the query command. Using bcftools (bcftools query -f ‘%POS\n’ chr21_10_5.bcf) it took 8m57s to extract the position values for the 2,365,367 variants from the BCF file. Using vcztools (vcztools query -f ‘%POS\n’ chr21_10_5.vcz) required 51s (note that this operation could be performed much more quickly and may be worth optimising for within vcztools, see Fig 5). This illustrates the general principle that vcztools only retrieves information necessary to fulfil a particular query, which can have a significant impact with large datasets.

To form a practical replacement for bcftools in production pipelines it is important that vcztools can produce VCF text at a reasonable rate. To do this we measure the time taken to generate VCF text for the last 10,000 variants in the dataset (using the previously extracted position column to determine the coordinates), vcztools view chr21_10_5.vcz -r1:47958263-. This took 30s, at a rate of 124.3 MiB of VCF text per second. The bcftools command on the corresponding BCF file took 25s (152.64 MiB/s) and on the VCF file it took 63s (60.57 MiB/s).

Vcztools is primarily intended to used as a replacement for bcftools in existing workflows based on pipes, where VCF text is sequentially filtered and processed in the classical Unix manner. In this context the most important property is that vcztools can produce VCF text as quickly as other processes in the pipeline can consume it. To get an estimate of the rate at which VCF can be parsed, we wrote a simple C program that reads a VCF file and parses it line-by-line using htslib [97]. For the 10,000 variant subset discussed above (3.73 GiB) this took 31s, giving a parsing rate of 123.1 MiB/s (almost identical to the write-rate of vcztools above). Given this limiting factor on overall throughput, it is unlikely that vcztools view would be a bottleneck in practical processing pipelines.

Vcztools is implemented in Python and C, and is available for download from the Python package index. Based on the region_index array computed by vcf2zarr, we use PyRanges [126] to perform efficient interval search queries.

### Interoperability: SAIGE proof-of-concept

To demonstrate the potential for integrating VCF Zarr into existing high-performance analysis tools, we developed a proof-of-concept extension of SAIGE [55, 56], a popular package for performing genome wide association tests. Using the TensorStore C++ library [103] we added support for VCF Zarr via an additional back-end (SAIGE reads data from PLINK, BGEN, VCF, BCF and SAV formats), for which the core functionality is defined in a 264 line C++ file. TensorStore provides efficient read and write access to array data in several formats, used, for example, in checkpointing [127] TensorFlow model training [128]. It supports several cloud platforms, and is based on an asynchronous, parallel processing model (see Discussion).

To assess the performance of this implementation we compared the time required to perform an association test using the simulated dataset of 100,000 samples from Fig 2. We performed a single-variant association test based on a phenotype simulated using tstrait [129], and measured the overall time required when using the equivalent VCF (162m), BCF (70m), SAV (7m) and Zarr (20m) files. Thus, in this case, substantially improved performance over the most popular formats, along with native cloud-store support are possible with relatively minor coding effort. We validated the implementation by comparing the association test results with the output of the other backends, confirming that they were numerically identical. As TensorStore does not currently support string data for Zarr, however, we were not able to fully replicate the output of fields such as MarkerID, Allele1 etc.

### Future applications: Cloud computing

To explore the use of cloud computing services with VCF Zarr, Genomics England provided access to aggV2 data in VCF Zarr format within their secure cloud environment. The VCF Zarr files for Chromosome 2 described in Table 1 were copied to a standard AWS S3 bucket. Using this data we repeated the benchmarks above on the same subset of 562,640 variants and 78,195 samples using the same single-threaded sequential Zarr code. Extracting the position array required 15s CPU, 23.8s elapsed. The af-dist calculation required 2.70min CPU, 5.23min elapsed. Creating the filter mask required 3.05min CPU, 11.15min elapsed.

While these benchmarks are comparable with results obtained on the shared file-system in the Genomics England research environment, the sequential single-threaded access patterns used do not map well to modern cloud computing environments. As each request to a cloud object store has a high latency, it is important to have many requests running concurrently to achieve good performance [100]. To explore the limits of performance we developed a simple prototype Zarr implementation, designed to maximise concurrency using asynchronous IO and multiple processes (see Methods for details). Fig S4 shows the rate at which genotypes can be decoded to RAM along with the rate at which we can compute af-dist (using Numba to classify genotypes, as before) as we increase the number of parallel processes on an EC2 c5.24xlarge instance with 96 vCPUs (48 physical). The maximum decompression rate of 47.4 GiB/s occurs at 64 processes (each with 100 concurrent requests), with a corresponding network data rate of 121 MiB/s. Fig S5 explores the limitations imposed by network access by measuring the rate at which we can download the chunks into RAM (without decompressing) using the same parallel and asynchronous architecture. Here, the maximum rate of 1.2 GiB/s for a c5.9xlarge instance and 2.4 GiB/s for a c5n.9xlarge (high performance network) instance was achieved with 32 processes. While these measurements are simplistic and may not be an accurate predictor of sustained bandwidth, it is still clear that genotype data can be fetched from S3 around an order of magnitude more quickly than it can be decompressed into RAM. We have also not measured the achievable bandwidth while scaling out across many instances, but given the ubiquity of cloud-based services and the scale at which they operate, it seems reasonable to assume that the claimed terabits per second of aggregate bandwidth from S3 [130] are achievable.

Returning to Fig S4, we can see that the af-dist calculation is performed at a maximum rate of 25.5 GiB/s using 128 processes. Genome wide, the aggV2 dataset consists of 722 million variants for 78,195 samples, giving a total of around 103 TiB of genotype data. We could therefore process the entire dataset at this maximum rate in about 69 minutes, with a compute cost of $5.55 (on-demand hourly rate of $4.848 for a c5.24xlarge instance in the eu-west-2 region, as-of 2025-01-31 [131]). AWS and other major clouds charge per object request (i.e. without considering the number of bytes transferred) and so the IO cost of this operation is determined by the number of chunks. As each 10,000 × 1,000 chunk encodes 19 MiB of genotype data, we incur about 5.7 million object requests to retrieve all chunks. At $0.00042 per 1000 GET requests for S3 Standard storage (eu-west-2 region as-of 2025-01-31 [132]), this comes to $2.39. Thus, it should be possible to perform af-dist and similar calculations genome wide on aggV2 for less than $10. Assuming 50KiB per chunk (the average stored chunk size for chromosome 2 is 47.57 KiB) we would need a total of 272 GiB to store the genotype data for the entire dataset. At $0.024 per GiB per month for S3 Standard (eu-west-2 region as-of 2025-01-31 [132]) this comes to a yearly storage cost of $78.33 (note that this is just the genotype data, not including other, much larger, fields listed in Table 1).

One of our benchmarks in the Genomics England case study section was to compare a straightforward sequential Zarr implementation of af-dist with bcftools on the genome slice corresponding to one of the original 106 VCF files for chromosome 2. There, Zarr’s efficient data format allows us to perform the same calculation in 4.25 minutes vs almost 12 hours for VCF. Here, by adapting our code to the high-throughput, high-variance nature of cloud stores and using the natural chunk-based parallelism of Zarr we reduce the elapsed time to 4.15 seconds. These benchmarks illustrate the transformative potential of Zarr and cloud computing.

### Future applications: GPU acceleration

GPU acceleration has revolutionised scientific computing by leveraging their massively parallel architecture to achieve significant speedups in computationally intensive tasks such as simulations, data analysis, and machine learning, often by one or two orders of magnitude [133, 134, 135] This has had a transformative impact across many fields [e.g. 136, 137]. While early GPU programming required expertise in low-level languages like CUDA or OpenCL, the advent of high-level Python libraries such as Numba [98], CuPy [138] and PyCUDA/PyOpenCL [139] has made GPU acceleration far more accessible. Deep learning [140] has led to major breakthroughs in several domains in recent years [e.g. 141, 142] and depends crucially on processing large volumes of data on GPUs or other custom hardware accelerators. The Python data science ecosystem is particularly powerful in this area, with GPU-native libraries like PyTorch [143] and TensorFlow [128] providing the basic infrastructure for the field.

To demonstrate the ease and efficiency with which data stored in Zarr format can be processed on a GPU, we developed two proof-of-concept applications. In both cases we demonstrate on data from chromosome 22 of 1000 Genomes WGS data [144]. This consists of 96,514 variants and 2504 samples, converted to Zarr format using a chunk size of 10,000 variants × 1,000 samples. Experiments in this section were performed on a computer with an Intel Xeon Silver 4216 processor, Nvidia A6000 RTX GPU and NVMe SSD.

To demonstrate the ease at which non-trivial, novel calculations can be performed on a GPU, we first reimplemented the af-dist calculation described above using CuPy [138], a GPU accelerated library of core operations for scientific computing. All of the operations required to compute af-dist are performed on the GPU, mostly with minimal changes required (e.g. replacing np.digitize with cp.digitize). We perform the central operation of genotype classification using the CuPy JIT compiler to generate a custom CUDA kernel. This is very similar to the Numba compiled function used in other experiments, with minimal changes required to take advantage of on-GPU parallelism. In the simplest version, we iterate over the chunks in the call_genotype array (stored as 30 chunks, consuming a total of 7.82 MiB), decompress each to a local NumPy array and then copy the data to GPU memory by creating the corresponding CuPy array. For this version, the total time in the genotype classification kernel was 25ms with a data transfer time of 179ms (we discount decompression time here; see next paragraph). For comparison, our standard Numba compiled classification function required a total of 1.46s running on a single CPU core. Another approach in which we load the entire call_genotype array into GPU memory at once (possible here as the 96514× 2504× 2 array requires 460 MiB) and uses a more sophisticated kernel in which we map GPU resources to subsets of the array in two dimensions, reduces transfer time to 138ms and processing time to 6ms.

Chunk decompression comprises a significant fraction of processing time (see Figs S1,S4) and host-to-GPU data transfer is often a bottleneck. It is therefore useful in some applications to directly decompress Zarr chunks into GPU memory. The KvikIO [145] Python package provides a GPU implementation of Zarr, such that both data transfers (file and network IO) and decompression are performed directly on the GPU. To explore the performance of the GPU optimised compression codecs available, we recompressed the call_AD (allele depth) array which had dimensions of 96514 × 2504 × 7 and 16 bit integer data type (note that we did not use local alleles encoding here, and this array is therefore highly sparse and compressible). For all five codecs considered (Bitcomp, Snappy, LZ4, Cascaded, Gdeflate) it took approximately 1s to compress and store this 3.15 GiB array. We then measured the time to decompress these arrays and copy back to host memory, which varied from 0.57s (Cascaded, LZ4) to 0.97s (Gdeflate) across the codecs. For reference, using the Zstandard codec and the standard CPU Zarr code (which performs some limited multithreading for chunk decompression by default) required about 7.6s.

These experiments are not formal benchmarks and are intended simply as illustative examples of the diverse, powerful and rapidly growing toolchain available with Zarr. Efficient access to VCF data in GPU memory may enable a number of downstream applications.

## Discussion

VCF is a central element of modern genomics, facilitating the exchange of data in a large ecosystem of interoperating tools. Its current row-oriented form, however, is fundamentally inefficient, profoundly limiting the scalability of the present generation of bioinformatics tools. Large scale VCF data cannot currently be processed without incurring a substantial economic (and environmental [146]) cost. We have shown here that this is not a necessary situation, and that greatly improved efficiency can be achieved by using more appropriate storage representations tuned to the realities of modern computing. We have argued that Zarr provides a powerful basis for cloud-based storage and analysis of large-scale genetic variation data. We propose the VCF Zarr specification which losslessly maps VCF data to Zarr, and provide the basic software infrastructure required for bidirectional conversion.

Zarr provides pragmatic solutions to some of the more pressing problems facing the analysis of large-scale genetic variation data, but it is not a panacea. Firstly, any dataset containing a variant with a large number of alleles (perhaps due to indels) will cause problems because the dimensions of fields are determined by their *maximum* dimension among all variants. In particular this is problematic for fields like PL in which the dimension depends quadratically on the number of alleles. While the local alleles representation [147] is a substantial improvement (see All of Us and Norway Spruce case studies) it may be lossy (see Methods). Secondly, the design of VCF Zarr emphasises efficiency of analysis for a fixed dataset, and does not consider how samples (and the corresponding novel variants) should be added. Thirdly, Zarr works best for numerical data of a fixed dimension, and therefore may not suitable for representing the unstructured data often included in VCF INFO fields.

Nonetheless, there are numerous datasets that exist today that would likely reap significant benefits from being deployed in a cloud-native fashion using Zarr. Object stores typically allow for individual objects (chunks, in Zarr) to be associated with “tags”, which can then be used to associate storage class, user access control and encryption keys. Aside from the performance benefits we have focused on here provided by Zarr, the ability to (for example) use high-performance storage for commonly used data such as the variant position and more cost-effective storage classes for infrequently used bulk QC data should provide significant operational benefits. Granular access controls would similarly allow non-identifiable variant-level data to be shared relatively freely, with genotype and other data more tightly controlled as required. Even finer granularity is possible if samples are grouped by access level within chunks (padding partially filled chunks as needed and using an appropriate sample mask). Providing client applications direct access to the data over HTTP and delegating access control to the cloud provider makes custom web APIs [148] and cryptographic container formats [149] largely unnecessary in this setting.

Modern computing is increasingly parallel, from multicore CPUs, GPUs and the massively parallel scale provided by public clouds. Making optimal use of these parallel resources requires new programming models, and in particular adapting analytics workloads to the properties of disaggregated cloud object stores is essential. Classical workflows in genomics emphasise streaming, in which large volumes of data are read sequentially from file, streamed through the processor, and the results written sequentially to storage or to the input of another process in a pipeline. Using this model on data held on a cloud object store requires us to emulate file-system semantics, and places hard requirements on receiving data in a specific order. As Durner et al [100] show (and we confirm in Fig S5), sustained data throughput speeds similar to a locally attached SSD can be obtained from cost-effective and massively scalable cloud object stores like Amazon S3, but on the condition that we do *not* place constraints on the order in which data is received (due to the high variance in latency). In this context, Zarr provides a simple and elegant model for computation in which the *chunk* is the basic unit of parallelism. For many applications, each chunk can be processed independently and in no particular order.

There is therefore an opportunity to develop a new generation of Zarr-native applications, taking advantage of the efficient data representation and natural model of chunk-based parallelism. Although there are several independent Zarr implementations across programming languages (see Methods), the Python data science ecosystem is a particularly powerful platform, with a rich suite of tools [e.g. 150, 151, 98, 152, 99, 153] and is increasingly popular in recent biological applications [e.g. 154, 155, 156, 157]. Xarray [158] provides a unified interface for working with multi-dimensional arrays in Python, and libraries like Dask [159] and Cubed [160] allow these operations to be scaled out transparently across processors and clusters. This scaling is achieved by distributing calculations over grid-based array representations like Zarr, where chunks provide the basic unit for parallel computation. The VCF Zarr specification introduced here was created to facilitate work on a scalable genetics toolkit for Python [161] built on Xarray. While the high-level facilities for distributed computation provided by Xarray are very powerful, they are not needed or indeed appropriate in all contexts. Our benchmarks here illustrate that working at the lowest level, by applying optimised kernels on a chunk-by-chunk basis is both straightforward to implement and highly performant. Thus, a range of possibilities exist in which developers can build utilities using the VCF Zarr specification using the appropriate level of abstraction and tool chain on a case-by-case basis.

While Zarr is now widely used across the sciences (see Methods) it was originally developed to store genetic variation data from the *Anopheles gambiae* 1000 Genomes Project [162] and is in active use in this setting [e.g. 163, 164]. The VCF Zarr specification presented here builds on this real-world experience but is still a draft proposal that would benefit from wider input across a range of applications. With some refinements and sufficient uptake it may be suitable for standardisation [2]. The benefits of Zarr are substantial, and, in certain settings, worth the cost of retooling away from classical file-oriented workflows. For example, the MalariaGEN Vector Observatory currently uses Zarr to store data from whole-genome sequencing of 23,000 *Anopheles* mosquitoes from 31 African countries [165]. The data is hosted in Google Cloud Storage and can be analysed interactively using free cloud computing services like Google Colab, enabling the use of data by scientists in malaria-endemic countries where access to local computing infrastructure and sufficient network bandwidth to download large datasets may be limited. VCF Zarr could similarly reduce the costs of analysing large-scale human data, and effectively open access to biobanks for a much broader group of researchers than currently possible.

## Methods

### Zarr and block-based compression

In the interest of completeness it is useful to provide a high-level overview of Zarr and the technologies that it depends upon. Zarr is a specialised format for storing large-scale *n*-dimensional data (arrays). Arrays are split into chunks, which are compressed and stored separately. Chunks are addressed by their indexes along the dimensions of the array, and the compressed data associated with this key. Chunks can be stored in individual files (as we do here), but a wide array of different storage backends are supported including cloud object stores and NoSQL databases; in principle, Zarr can store data in any key-value store. Metadata describing the array and its properties is then stored in JSON format along with the chunks. The simplicity and transparency of this design has substantial advantages over technologies such as HDF [166] and NetCDF [167] which are based on complex layouts of multidimensional data within a single file, and cannot be accessed in practice without the corresponding library. (See [41] for further discussion of the benefits of Zarr over these monolithic file-oriented formats.) In contrast, there are numerous implementations of the Zarr specification, ranging from the mature Zarr-Python [101] and TensorStore [103] implementations to more experimental extensions to packages like GDAL [168], NetCDF [169], N5 [170] and xtensor [171] as well as standalone libraries for C++ [172], Java [102], JavaScript [173], Julia [174], Rust [175] and R [176].

Zarr is flexible in allowing different compression codecs and pre-compression filters to be specified on a per-array basis. Two key technologies often used in conjunction with Zarr are the Blosc meta-compressor [95] and Zstandard compression algorithm [94]. Blosc is a high-performance compressor optimised for numerical data which uses “blocking” [95] to optimise CPU-cache access patterns, as well as highly optimised bit-wise and byte-wise shuffle filters. Remarkably, on highly compressible datasets, Blosc decompression can be faster than memcpy. Blosc is written in C, with APIs for C, Python, Julia, Rust and others. Blosc is a “meta-compressor” because it provides access to several different compression codecs. The Zstandard codec is of particular interest here as it achieves very high compression ratios with good decompression speeds (Figs S1, S7). Zstandard is also used in several recent VCF compression methods [e.g. 65, 66].

Scientific datasets are increasingly overwhelming the classical model of downloading and analysing locally, and are migrating to centralised cloud repositories [41, 177]. The combination of Zarr’s simple and cloud-friendly storage of data chunks with state-of-theart compression methods has led to Zarr gaining significant traction in these settings (see [178, 177, 179] Zarr cloud benchmarks). Multiple petabyte-scale datasets are now stored using Zarr [e.g. 179, 180, 181] or under active consideration for migration [178, 182]. The Open GeoSpatial consortium has formally recognised Zarr as a community standard [183] and has formed a new GeoZarr Standards Working Group to establish a Zarr encoding for geospatial data [184].

Zarr has recently been gaining popularity in biological applications. The Open Microscopy Environment has developed OME-Zarr [185] as one of its “next generation” cloud ready file formats [177]. OME-Zarr already has a rich suite of supporting tools [185, 186]. Zarr has also seen recent uptake in other large-scale processing for microscopy applications [172], as well as single-cell genomics [187, 188] and multimodal spatial omics data [189, 190]. Recent additions using Zarr include the application of deep learning models to genomic sequence data [191], storage and manipulation of large-scale linkage disequilibrium matrices [192], and a browser for genetic variation data [193].

### The VCF Zarr specification

The VCF Zarr specification is a direct mapping from the VCF data model to a chunked binary array format using Zarr, and is an evolution of the Zarr format used in the scikit-allel package [194]. VCF Zarr takes advantage of Zarr’s hierarchical structure by representing a VCF file as a top-level Zarr group containing Zarr arrays. Each VCF field (fixed fields, INFO fields, and FORMAT fields) is represented as a separate array in the Zarr hierarchy. Some of the structures from the VCF header are also represented as arrays, including contigs, filters, and samples.

The specification defines the name, shape, dimension names, and data type for each array in the Zarr store. These “logical” properties are mandated, in contrast to “physical” Zarr array properties such as chunk sizes and compression, which can be freely chosen by the implementation. This separation makes it straightforward for tools and applications to consume VCF Zarr data since the data has a well-defined structure, while allowing implementations enough room to optimise chunk sizes and compression according to the application’s needs.

The specification defines a clear mapping of VCF field names (keys) to array names, VCF Number to array shape, and VCF Type to array data type. To take one example, consider the VCF AD genotype field defined by the following VCF header: ##FORMAT=<ID=AD,Number=A,Type=Integer,Description=“Allele Depths”>. The FORMAT key ID maps to an array name of call_AD (FORMAT fields have a call_ prefix, while INFO fields have a variant_ prefix; both are followed by the key name). Arrays corresponding to FORMAT fields are 3-dimensional with shapes that look like (variants, samples, <Number>) in general. In this case, the Number A entry indicates that the field has one value per alternate allele, which in VCF Zarr is represented as the alt_alleles dimension name, so the shape of this array is (variants, samples, alt_alleles). The VCF Integer type can be represented as any Zarr integer type, and the specification doesn’t mandate particular integer widths. The vcf2zarr (see the next section) conversion utility chooses the narrowest integer width that can represent the data in each field.

An important aspect of VCF Zarr is that field dimensions are global and fixed, and defined as the maximum across all rows. Continuing the example above, the third dimension of the array is the maximum number of alternate alleles across *all* variants. For variants at which there are less than the maximum number of alternative alleles, the third dimension of the call_AD array is padded with a sentinel value (−2 for integers and a specific non-signalling NaN for floats). This is problematic in situations where some sites have a large number of alleles (e.g., 95 in the All of Us case study above), and in particular for fields that have a quadratic dependency on the number of alleles (Number=G) such as PL. We use the local alleles encoding [147] to reduce the storage requirements for such fields (see the next section for implementation details).

The VCF Zarr specification can represent anything described by BCF (which is somewhat more restrictive than VCF) except for two corner cases related to the encoding of missing data. Firstly, VCF Zarr does not distinguish between a field that is not present and one that is present but contains missing data. For example, a variant with an INFO field NS=. is represented in the same way in VCF Zarr as an INFO field with no NS key. Secondly, because of the use of sentinel values to represent missing and fill values for integers (−1 and -2, respectively), a field containing these original values cannot be stored. In practice this doesn’t seem to be much of an issue (we have not found a real VCF that contains negative integers). However, if -1 and -2 need to be stored, a float field can be used without issues.

The VCF Zarr specification is general and can be mapped to file formats such as PLINK [19, 20] and BGEN [21] with some minor extensions.

### vcf2zarr

Converting VCF to Zarr at Biobank scale is challenging. One problem is to determine the dimension of fields, (i.e., finding the maximum number of alternate alleles and the maximum size of Number=. fields) which requires a full pass through the data. Another challenge is to keep memory usage within reasonable limits: although we can view each record in the VCF one-by-one, we must buffer a full chunk (10,000 variants is the default in vcf2zarr) in the variants dimension for each of the fields to convert to Zarr. For VCFs with many FORMAT fields and large numbers of samples this can require tens of gigabytes of RAM per worker, making parallelism difficult. Reading the VCF multiple times for different fields is possible, but would be prohibitively slow for multi-terabyte VCFs.

The vcf2zarr utility solves this problem by first converting the VCF data (which can be split across many files) into an Intermediate Columnar Format (ICF). The vcf2zarr explode command takes a set of VCFs, and reads through them using cyvcf2 [195], storing each field independently in (approximately) fixed-size compressed chunks in a file-system hierarchy. ICF is designed to support efficient Zarr encoding within vcf2zarr and not intended for reuse outside that context. Large files can be partitioned based on information extracted from the CSI or Tabix indexes, and so different parts of a file can be converted to ICF in parallel. Once all partitions have completed, information about the number of records in each partition and chunk of a given field is stored so that the record at a particular index can be efficiently retrieved. Summaries such as maximum dimension and the minimum and maximum value of each field are also maintained, to aid choice of data types later. A set of VCF files can be converted to intermediate columnar format in parallel on a single machine using the explode command, or can be distributed across a cluster using the dexplode-init, dexplode-partition and dexplode-finalise commands.

Once the VCF data has been converted to the intermediate columnar format, it can then be converted to Zarr using the vcf2zarr encode command. By default we choose integer widths based on the maximum and minimum values observed during conversion to ICF along with reasonable compressor defaults (see next section). Default choices can be modified by generating a JSON-formatted storage schema, which can be edited and supplied as an argument to encode. Encoding a given field (for example, call_AD) involves creating a buffer to hold a full variant-chunk of the array in question, and then sequentially filling this buffer with values read from ICF and flushing to file. Similar to the explode command, encoding to Zarr can be done in parallel on a single machine using the encode command, or can be distributed across a cluster using the dencode-init, dencode-partition and dencode-finalise commands. The distributed commands are fault-tolerant, reporting any failed partitions so that they can be retried.

The implementation of the local alleles data reduction technique in vcf2zarr follows the approach of Poterba et al [147]. Specifically, we identify the alleles in use for each call by examining the genotypes and then store the LA field, listing the (maximum of ploidy) unique alleles observed as indexes into the global list for that variant. Using the LA field we then compute derived fields such as AD and PL in terms of the local alleles during Zarr encoding (see e.g. the call_LAD field in Table 3 and call_LPL field in Table 4). Note that our approach (along with Poterba et al) differs from the details in the VCF 4.5 specification [196], which defines the LAA field containing indexes of ALT alleles only. Explicitly listing the alleles observed leads to simpler code and means we do not need to refer back to the genotypes to distinguish missing data from homozygous reference calls. Note also that the local alleles encoding may be lossy, as only information relevant to the chosen alleles for each call is retained.

To facilitate efficient region search queries, we compute an additional array region_index which stores information about the range of genomic positions in each chunk and provides sufficient information to implement interval search queries. We can then load this index into memory and use it to determine which variant chunks need to be retrieved. In the Norway Spruce example (Table 4), which has 3.75 billion variants, this index requires 3.14 MiB of storage over 12 chunk files to store information about all 374,518 variant chunks (the variant_position array, for comparison is 13.95 GiB uncompressed).

### Choosing default compressor settings

To inform the choice of compression settings across different fields in VCF data, we analysed their effect on compression ratio on recent high-coverage WGS data from the 1000 Genomes project [144]. We began by downloading the first 100,000 lines of the VCF for chromosome 22 (giving a 1.1 GiB compressed VCF) and converted to Zarr using vcf2zarr with default settings. We then systematically examined the effects of varying chunk sizes and compressor settings on the compression ratio for call-level fields. (We excluded call_PL from this analysis as this analysis predated our implementation of local alleles.)

Fig S7 shows the effect of varying compression codecs in Blosc. The combination of outstanding compression performance and competitive decoding speed (Fig S1) makes zstd a good default choice.

The shuffle parameter in the Blosc meta-compressor [95] can result in substantially better compression, albeit at the cost of somewhat slower decoding (see Fig S1). Fig S8 shows the effect of bit shuffle (grouping together bits at the same position across bytes before compression), and byte shuffle (grouping together bytes at the sample position across words before compression) on compression ratio. Bit shuffle provides a significant improvement in compression for the call_genotype field because the vast majority of genotype calls will be 0 or 1, and therefore bits 1 to 7 will be 0. Thus, grouping these bits together will lead to significantly better compression. This strategy also works well when compressing boolean fields stored as 8 bit integers, where the top 7 bits are always 0. In practice, boolean fields stored in this way have very similar compression to using a bit-packing pre-compression filter (data not shown). Although byte shuffle leads to somewhat better compression for call_AD and call_DP, it gives substantially worse compression on call_AB than no shuffling. The default in vcf2zarr is therefore to use bit shuffle for call_genotype and all boolean fields, and to not use byte shuffling on any field. These defaults can be easily overruled, however, by outputting and modifying a JSON formatted storage schema before encoding to Zarr.

Fig S9 shows that chunk size has a weak influence on compression ratio for most fields. Increasing sample chunk size slightly increases compression on call_AB, and has no effect on less compressible fields. Variant chunk size appears to have almost no effect on compression ratio. Interestingly, the choice of chunk size along the sample dimension for the genotype matrix does have a significant effect. With six evenly spaced points between 100 and 2504, Fig S9A shows a somewhat unpredictable relationship between sample chunk size and compression ratio. The more fine-grained analysis of Fig S10 shows that three distinct trend lines emerge depending on the chunk size divisibility, with the modulus (i.e., the remainder in the last chunk) also having a minor effect. At greater than 40X, compression ratio is high in all cases, and given that genotypes contribute relatively little to the total storage of real datasets (Table 1) the effect will likely be fairly minor in practice. Thus, we do not expect the choice of chunk size to have a significant impact on overall storage usage, and so choice may be determined by other considerations such as expected data access patterns.

### Benchmarks

In this section we describe the methodology used for the simulation-based benchmarks of Figs 2,3, 4 and 5. The benchmarks use data simulated by conditioning on a large pedigree of French-Canadians using msprime [197], which have been shown to follow patterns observed in real data from the same population to a remarkable degree [93]. We begin by downloading the simulated ancestral recombination graph [198, 199, 200] for chromosome 21 from Zenodo [201] in compressed tszip format. This 552M file contains the simulated ancestry and mutations for 1.4 million present-day samples. We then subset the full simulation down to 10^1^, 10^2^, …, 10^6^ samples using ARG simplification [202, 200], storing the subsets in tskit format [203, 204]. Note that this procedure captures the growth in the number of variants (shown in the top x-axis labels) as we increase sample sizes as a natural consequence of population-genetic processes. As a result of simulated mutational processes, most sites have one alternate allele, with 7.9% having two and 0.2% having three alternate alleles in the 10^6^ samples dataset. We then export the variation data from each subset to VCF using tskit vcf subset.ts | bgzip > subset.vcf.gz as the starting point for other tools.

We used bcftools version 1.18, Savvy 2.1.0, Genozip 5.0.26, vcf2zarr 0.0.9, and Zarr-Python 2.17.2. All tools used default settings, unless otherwise stated. All simulation-based benchmarks (excluding Fig S2) were performed on a dual CPU (Intel Xeon E5-2680 v2) server with 256 GiB of RAM running Debian GNU/Linux 11. To ensure that the true effects of having data distributed over a large number of files were reported, benchmarks for Zarr and Savvy were performed on a cold disk cache by running echo 3 | sudo tee /proc/sys/vm/drop_caches before each run. The I/O subsystem used is based on a RAID 5 of 12 SATA hard drives. For the CPU time benchmarks we measure the sum of the total user and system times required to execute the full command (as reported by GNU time) as well as elapsed wall-clock time. Total CPU time is shown as a solid line, with wall-clock time as a dashed line of the same colour. In the case of pipelines, where some processing is conducted concurrently wall-clock time can be less than total CPU (e.g. genozip in Fig 3). When I/0 costs are significant, wall-clock time can be greater than total CPU (e.g. Zarr and Savvy in Fig 4). Each tool was instructed to use one thread, where the options were provided. Where possible in pipelines we use uncompressed BCF output (-Ou) to make processing more efficient [54]. We do not use BCF output in genozip because it is not supported directly.

Because bcftools +af-dist requires the AF INFO field and this is not kept in sync by bcftools view (although the AC and AN fields are), the subset calculation for Fig 4 requires an additional step. The resulting pipeline is bcftools view -r REGION -S SAMPLESFILE -IOu BCFFILE | bcftools +fill-tags -Ou | bcftools +af-dist. Genozip similarly requires a +fill-tags step in the pipeline.

### AWS parallel Zarr prototype

This section describes the methodology used for the AWS parallel Zarr throughput measurements in Fig S4 and Fig S5, and timing measurements detailed in the cloud computing results. These measurements were run on EC2 instances using Jupyter notebooks hosted on Genomics England’s Lifebit CloudOS platform. VCF Zarr chunks were hosted in a standard profile S3 bucket. Both instance and bucket were in the same availability zone in the eu-west-2 region. Each of the parallel Python processes used the aioboto3 library (version 13.2.0) to fetch chunks concurrently from S3. To measure throughput accurately, the number of chunks fetched was scaled with the number of processes such that each process fetched 60 GiB of decompressed data. For decompressing and computing afdist the same architecture was used, but as fetched chunks became available in memory in each process, they were decompressed and processed synchronously. Decompression was performed by using the numcodecs library [112].

This prototype did not use any existing Zarr implementation as the levels of concurrency involved require specialised asynchronous programming techniques that are not widely supported. The fact that such a high-performance prototype can be implemented in a notebook with no Zarr-specific library support is a testament to the transparent simplicity of the Zarr format.

## Availability of source code and requirements

The VCF Zarr specification is available on GitHub at https://github.com/sgkit-dev/vcf-zarr-spec/. The SAIGE prototype is available at https://github.com/Will-Tyler/SAIGE and all other code for running benchmarks, analyses and creating plots in this article is available at https://github.com/sgkit-dev/vcf-zarrpublication. Vcf2zarr is freely available under the terms of the Apache 2.0 license as part of the bio2zarr suite (https://github.com/sgkit-dev/bio2zarr/) and can be installed from the Python Package Index (https://pypi.org/project/bio2zarr/). Vcztools (https://github.com/sgkit-dev/vcztools/) is freely available under the terms of the Apache 2.0 license and can be installed from the Python Package Index (https://pypi.org/project/vcztools/).

## List of abbreviations

ICF: Intermediate Columnar Format
GWAS: Genome Wide Association Study
PBWT: Positional Burrows-Wheeler Transform
QC: Quality Control
UKB: UK Biobank
VCF: Variant Call Format
WGS: Whole Genome Sequence

## Acknowledgements

We are grateful to Augusto Rendon and the Genomics England support team for facilitating our cloud computing experiments, and to Gil McVean and Ben Neale for helpful discussions and comments on the manuscript.

This research was made possible through access to data in the National Genomic Research Library, which is managed by Genomics England Limited (a wholly owned company of the Department of Health and Social Care). The National Genomic Research Library holds data provided by patients and collected by the NHS as part of their care and data collected as part of their participation in research. The National Genomic Research Library is funded by the National Institute for Health Research and NHS England. The Wellcome Trust, Cancer Research UK and the Medical Research Council have also funded research infrastructure.

This study makes use of de-identified data held by Our Future Health. We would like to acknowledge all the research participants who have donated their data to the Our Future Health research programme [205].

Computation used the Oxford Biomedical Research Computing (BMRC) facility, a joint development between the Wellcome Centre for Human Genetics and the Big Data Institute supported by Health Data Research UK and the NIHR Oxford Biomedical Research Centre. The views expressed are those of the author(s) and not necessarily those of the NHS, the NIHR or the Department of Health.

Computation for the Spruce case study were enabled by resources provided by the National Academic Infrastructure for Supercomputing in Sweden (NAISS), partially funded by the Swedish Research Council through grant agreement no. 2022-06725.

Genozip was used under the terms of the free Genozip Academic license. Genozip was only used on simulated data, in compliance with the “No Commercial Data” criterion.

## Supplementary Material

**Figure S1.**
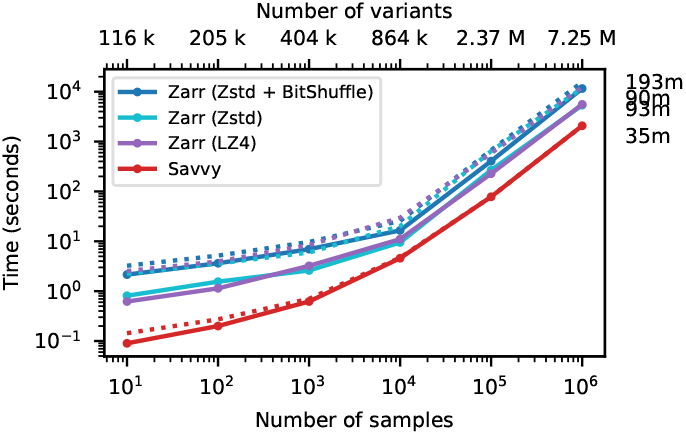
Genotype decoding performance. Total CPU time required to decode genotypes into memory using the Zarr-Python and Savvy C++ APIs for the data in Figure 2. Elapsed time is also reported (dotted line). This corresponds to a maximum rate of 1.2 GiB/s for Zarr (Zstd + BitShuffle), 2.7 GiB/s Zarr (Zstd), 2.9 GiB/s Zarr (LZ4), and 6.6 GiB/s for Savvy.

**Figure S2.**
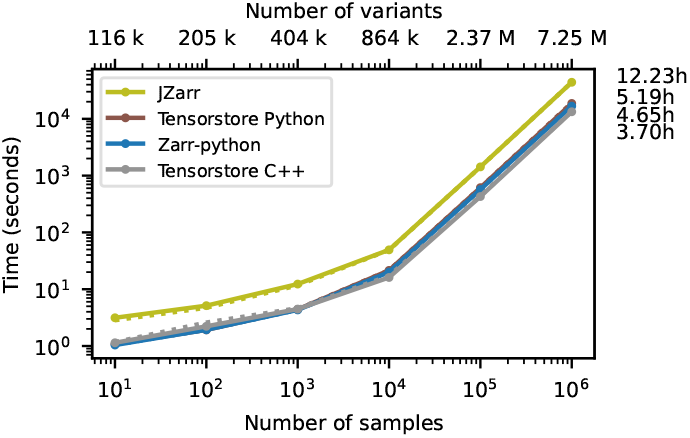
Whole-matrix computation performance using different Zarr implementations. Total CPU time required to run the af-dist calculation the data in Figure 2 using different Zarr implementations. Elapsed time is also reported (dotted line). These benchmarks were run on an 8-core CPU (Intel i7-9700) with 32 GiB RAM running Linux Mint 21.3 with data on an NVMe SSD. Note the difference between the time reported here and in Fig 3 for Zarr-Python is due to different hardware platforms.

**Figure S3.**
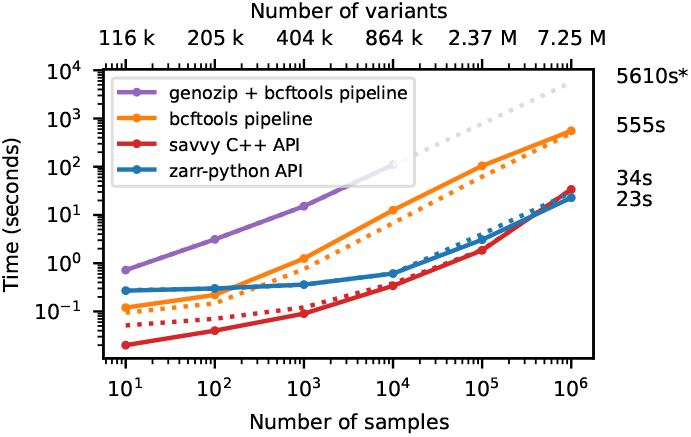
Compute performance on a large subset of the genotype matrix. Total CPU time required to run the af-dist calculation for a subset of half of the samples and 10000 variants from the middle of the matrix for the data in Figure 2. Elapsed time is also reported (dotted line). Genozip did not run for *n* > 10^4^ samples because it does not support a file to specify sample IDs, and the command line was therefore too long for the shell to execute.

**Figure S4.**
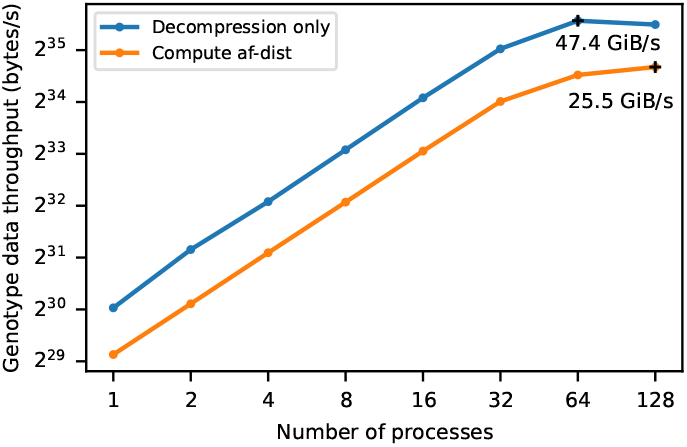
Scalability of genotype data processing on AWS. Using the prototype parallel Zarr implementation (see Methods) we measured the rate genotype data can be decoded to memory (decompression only) and the rate at which we can perform the full af-dist calculation as we vary the number of parallel processes.

**Figure S5.**
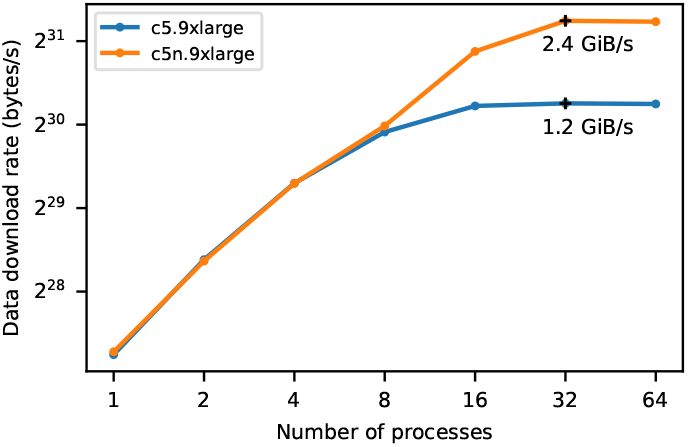
Chunk data download rate on AWS. Using the prototype parallel Zarr implementation (see Methods) we measured the rate at which chunks of compressed genotype data can be downloaded on two different instance types.

**Figure S6.**
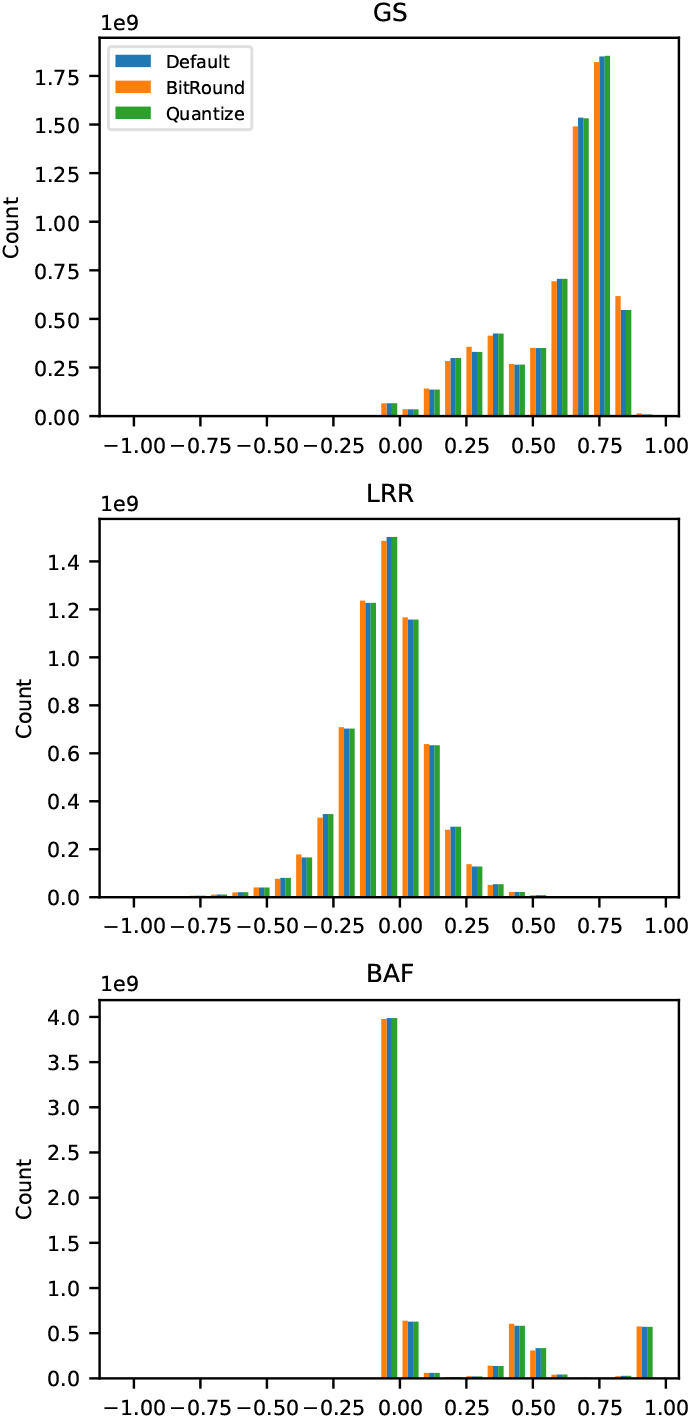
Distribution of values in the GS, LRR and BAF fields in the Our Future Health data with no truncation (Default), and truncation using the BitRound and Quantize filters.

**Figure S7.**
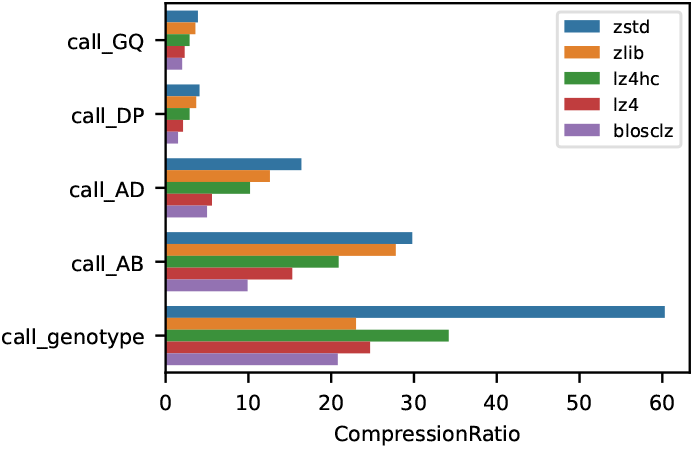
Effects of Blosc compression codec on compression ratio on call-level fields in 1000 Genomes data. In all cases compression level=7 was used, with a variant chunk size of 10,000 and sample chunk size of 1,000. Bit shuffle was used for call_genotype, and no shuffle used for the other fields.

**Figure S8.**
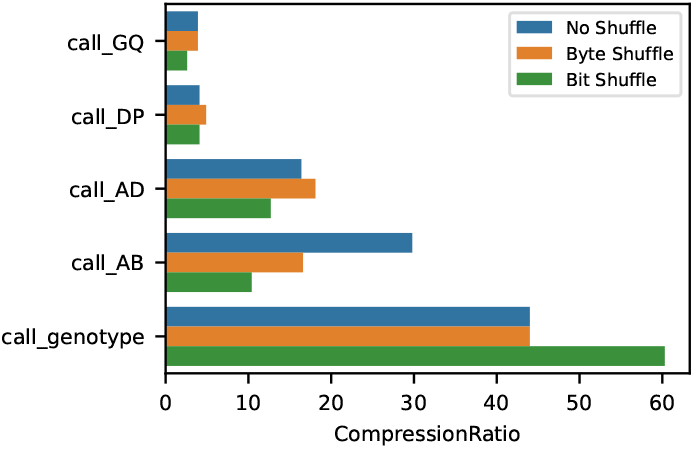
Effects of Blosc shuffle settings on compression ratio on call-level fields in 1000 Genomes data. In all cases the zstd compressor with compression level=7 was used, with a variant chunk size of 10,000 and sample chunk size of 1,000.

**Figure S9.**
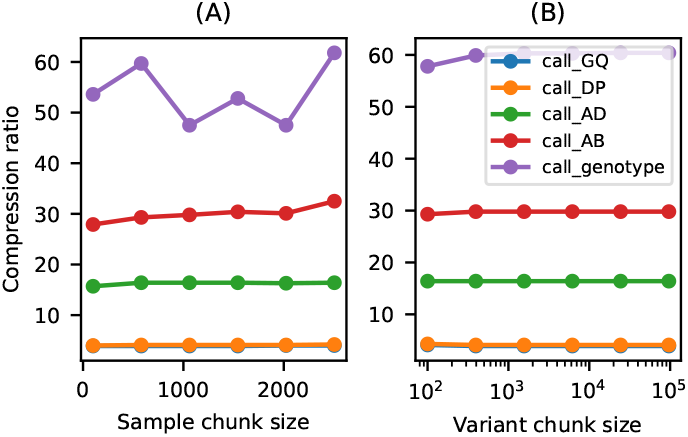
Effects of chunk sizes on compression ratio on call-level fields in 1000 Genomes data. (A) Varying sample chunk size, holding variant chunk size fixed at 10,000. (B) Varying variant chunk size, holding sample chunk size fixed at 1,000. In all cases the zstd compressor with compression level=7 was used. Bit shuffle was used for call_genotype, and no shuffle used for the other fields. Values are chosen to be evenly spaced on a linear scale between 100 and 2504 (the number of samples) in and evenly spaced between 100 and 96514 on a log scale in (B).

**Figure S10.**
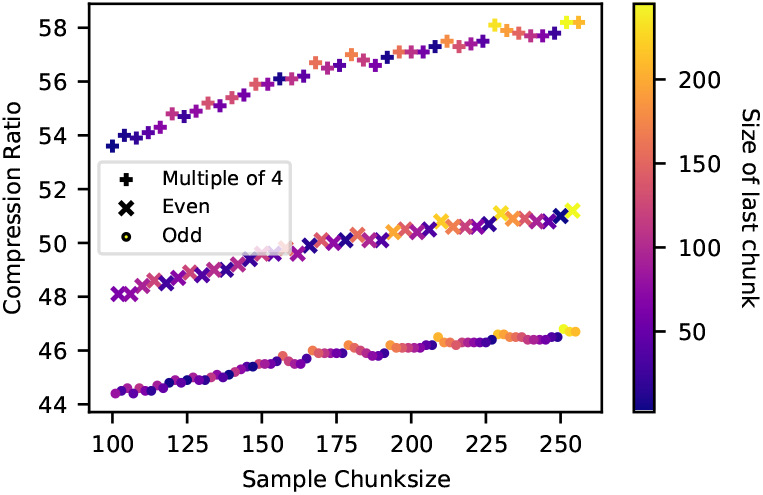
Effects of sample chunk size on compression ratio on the call_genotype field in 1000 Genomes data. The same analysis as in Fig S9, except we only consider call_genotype and we examine all sample chunk sizes from 100 to 256. Distinct trendlines emerge for odd, even and multiple-of-four chunk sizes (shown by markers). The size of the final chunk also has a minor effect (shown by colour).

